# Metabolomic rearrangement controls the intrinsic microbial response to temperature changes

**DOI:** 10.1101/2023.07.22.550177

**Authors:** Benjamin D. Knapp, Lisa Willis, Carlos Gonzalez, Harsh Vashistha, Joanna Jammal Touma, Mikhail Tikhonov, Jeffrey Ram, Hanna Salman, Josh E. Elias, Kerwyn Casey Huang

## Abstract

Temperature is one of the key determinants of microbial behavior and survival, whose impact is typically studied under heat- or cold-shock conditions that elicit specific regulation to combat lethal stress. At intermediate temperatures, cellular growth rate varies according to the Arrhenius law of thermodynamics without stress responses, a behavior whose origins have not yet been elucidated. Using single-cell microscopy during temperature perturbations, we show that bacteria exhibit a highly conserved, gradual response to temperature upshifts with a time scale of ∼1.5 doublings at the higher temperature, regardless of initial/final temperature or nutrient source. We find that this behavior is coupled to a temperature memory, which we rule out as being neither transcriptional, translational, nor membrane dependent. Instead, we demonstrate that an autocatalytic enzyme network incorporating temperature-sensitive Michaelis-Menten kinetics recapitulates all temperature-shift dynamics through metabolome rearrangement, which encodes a temperature memory and successfully predicts alterations in the upshift response observed under simple-sugar, low-nutrient conditions, and in fungi. This model also provides a mechanistic framework for both Arrhenius-dependent growth and the classical Monod Equation through temperature-dependent metabolite flux.

## Introduction

To proliferate and survive, microbes must integrate a vast array of environmental signals such as pH^1^, salt^2^, osmolarity^3^, and nutrient availability^4^, whose effects are monitored and integrated through intracellular regulation. By contrast, while growth is critically dependent on the environmental temperature, microbes are unable to regulate intracellular temperature. Enteric bacteria face temperature changes across time scales ranging from minutes (host colonization) to hours or days (fever), and soil and marine species are regularly exposed to daily and seasonal fluctuations. The effects of temperature can be dramatic: the largest determinant of marine bacterial community composition is local sea temperature^5^, and long-term temperature changes associated with climate change are thought to substantially impact soil microbial biomass^6^. Despite these fundamental connections, the mechanistic underpinnings of how temperature affects microbial behavior remain unclear.

Most studies on the cellular effects of temperature have focused on the regulatory systems that enable survival under heat and cold stress, during which molecular chaperones assist in the folding and unfolding of proteins and RNA^7,8^. While growth rate decreases at extreme, stress response-inducing temperatures^9^, there is typically a temperature range over which growth rate increases with temperature in agreement with the Arrhenius Law of equilibrium thermodynamics^10,11^, wherein the natural logarithm of growth rate depends approximately linearly on the inverse of the absolute temperature, with a negative slope that can be interpreted as an activation energy for growth (*E_a_*, equivalent to the barrier for enzyme kinetics)^12^. This temperature-dependent behavior is highly conserved across diverse bacteria, archaea, yeast, and mammalian cells^12^, with each species exhibiting its own Arrhenius temperature range and activation energy.

While theoretical work has suggested that proteome stability sets the upper bound for the growth rate of a given bacterial species across temperatures^13^, there is no correlation between permissible (optimal) growth temperatures and *E_a_* across bacteria^10^, indicating that *E_a_* is likely influenced by factors other than proteome stability. In *Escherichia coli*, the Arrhenius range is between 23 °C and 37 °C and *E_a_* is ∼13 kcal/mol^11,13^, similar to the free energy change from ATP hydrolysis (12–16 kcal/mol)^14^, suggesting that there may be a single rate-limiting enzyme for growth. On the other hand, theoretical work on cyclical enzyme networks has suggested that *E_a_* arises from the average of the activation energies over all reactions within the network^15,16^, as the *E_a_* for most biological enzymes is constrained between 5–20 kcal/mol^12,17,18^. However, the factor(s) that limits growth across Arrhenius temperatures and sets *E_a_* for a given species remains unknown.

Most previous studies of growth in the Arrhenius range have focused on steady-state growth, rather than how cells respond to temperature changes. An early study of 133 proteins in *E. coli* showed that the concentration of most of these proteins was maintained across Arrhenius temperatures, suggesting that the proteome may be largely temperature insensitive^11^. However, other studies found transient changes in the synthesis of tRNA synthetases after temperature increases^19^, and a decrease from 37 °C to 28 °C resulted in significant changes to ∼9% of the *E. coli* transcriptome^20^, perhaps reflecting the selection for coupled sensing of changes in temperature and other environmental variables such as oxygen levels relevant for animal host colonization^21^. In the few studies examining the growth rate response to temperature shifts, some report that *E. coli* growth rate responds almost immediately upon temperature shifts within the Arrhenius range^19,22^, while others report much longer time scales that scale with the final temperature^23^.

Many factors could determine the time scale of temperature adaptation. *E. coli* cells tightly maintain membrane fluidity across all temperatures permissible for growth by regulating membrane composition^24^. Increased membrane fluidity allows for higher respiratory metabolic rates^25^, but how fluidity impacts growth rate during temperature shifts is unknown. Growth rate across nutrients correlates with ribosome concentration, which is optimized through competition between protein and autocatalytic ribosome synthesis^26,27^. As a result, many models of growth focus on translation as a key growth-limiting factor^28^. However, this framework is likely inappropriate for understanding growth rate variations across temperatures, as ribosome concentration is constant across Arrhenius temperatures^11,29^.

Here, we demonstrate that *E. coli* cells exhibit an asymmetric growth-rate response to temperature shifts within the Arrhenius range: during downshifts, growth-rate rapidly decreases on the same time scale as that of the temperature shift, while cells gradually adapt to upshifts on a time scale corresponding to ∼1.5 doublings at the steady-state growth rate of the final temperature, independent of nutrient source. We show that these responses do not result from proteome or membrane reconfiguration or from transcriptional regulation, although increased membrane fluidity was lethal at high temperatures. To interrogate the mechanism underlying growth-rate adaptation to temperature shifts, we developed an autocatalytic enzyme network model that incorporates temperature-sensitive Michaelis-Menten kinetics into chained reactions. The model quantitatively captured the asymmetric responses to up- and downshifts across Arrhenius temperatures and how carbon source affects the dynamics immediately after an upshift through changes in the activation energy of growth. Furthermore, our model successfully predicted the larger initial acceleration and decrease in upshift response time observed at sub-saturating substrate concentration compared with saturation. Growth-rate dynamics during temperature upshifts were highly conserved across diverse *Escherichia* strains/species and the Gram-positive *Bacillus subtilis*, suggesting that certain aspects of microbial responses to temperature shifts and hence features of the underlying metabolic network may be universally conserved. Moreover, the ability of the model to capture the distinct response of the fission yeast *Schizosaccharomyces pombe* suggests that temperature adaptation provides a window into the factors limiting growth.

## Results

### The response time to temperature upshifts is determined by the steady-state growth rate at the higher temperature

To determine whether evolutionary history impacts temperature sensitivity across strains of a single species, we measured the growth of 12 *E. coli* strains isolated from fecal samples of hosts encompassing a broad range of body temperatures (Table S1)^30,31^. All strains exhibited similar temperature sensitivity profiles, with Arrhenius behavior (linear relationship between ln(growth rate) and 1/T) between 27 °C and 37 °C (Fig. S1B), with similar activation energies (10–15 kcal/mol) of growth (Fig. S1C–E), consistent with a null model in which growth is determined by a single rate-limiting enzyme. Single-cell growth rates were highly correlated with bulk growth rates from liquid culture across the Arrhenius range (Fig. 1A, inset, S2). These data suggest that the thermodynamic properties of *E. coli* growth are conserved and that host body temperature does not dictate the temperature sensitivity of enteric bacteria.

**Figure 1:**
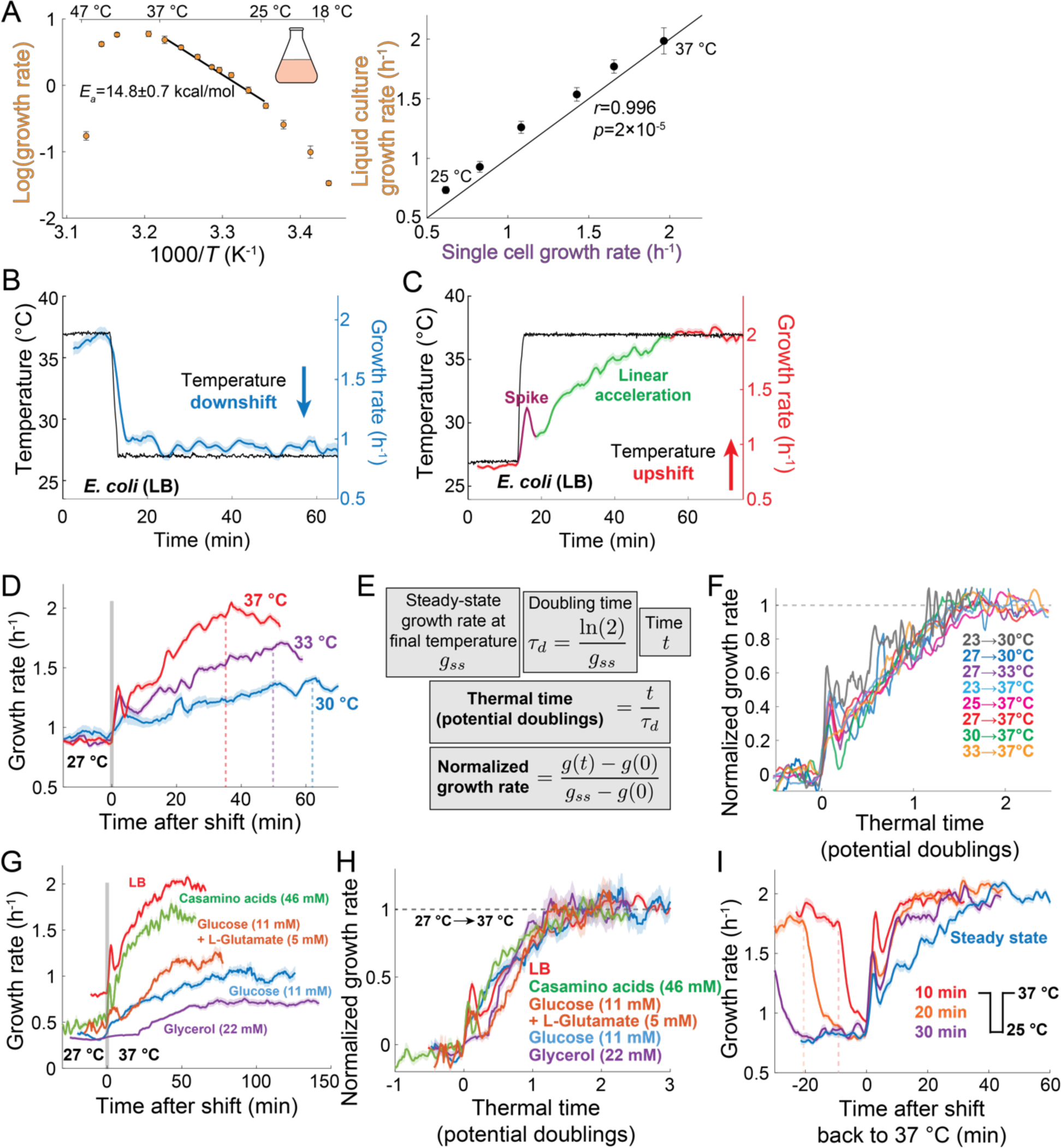
*E. coli* responds asymmetrically to temperature shifts, and upshift dynamics are determined by the steady-state growth rate at the final temperature. A) (Left) Arrhenius plot illustrating that the natural logarithm of *E. coli* MG1655 maximal growth rate for temperatures between 25 °C and 37 °C varies linearly with the inverse absolute temperature, with a negative slope whose magnitude is the activation energy (E_a_). (Right) Liquid-culture and single-cell growth rates were highly correlated for temperatures in the Arrhenius range between 25 °C and 37 °C (data from Fig. S1C,E). B) Single-cell growth rate (blue) of *E. coli* MG1655 throughout a temperature (black) downshift from 37 °C to 27 °C in rich growth medium (LB) (*n*=253 cells). The curve is the mean growth rate and the shaded region represents one standard error of the mean (SEM) Growth rate responds quickly to the downshift. C) Single-cell growth rate (red) of *E. coli* MG1655 throughout a temperature (black) upshift from 27 °C to 37 °C in LB (*n*=792 cells). The curve is the mean growth rate and the shaded regions represent ±1 SEM. Growth rate initially spikes (dark red), followed by linear acceleration (green) to the new steady-state value over ∼35 min. D) Single-cell growth rate of *E. coli* MG1655 throughout a temperature upshift from 27 °C to 30 °C (blue, *n*=253 cells), 33 °C (purple, *n*=754 cells), or 37 °C (red, *n*=904 cells) in LB. Vertical dashed lines indicate the time at which the growth rate reached its steady-state value at the higher temperature. Curves show the mean growth rate and the shaded regions represent ±1 SEM. E) The thermal time is defined as the time (*t*) measured in units of doubling times (*ρ*_d_) of steady-state growth at the higher temperature. The normalized growth rate is defined as the difference between the growth rate at time (*g*(*t*)) and the growth rate at the temperature shift (*g*(0)) divided by the difference between the steady-state growth rate at the higher temperature (*g_ss_*) and the initial temperature (*g*(0)). F) Normalized single-cell growth rate of *E. coli* MG1655 grown in LB exhibits a characteristic response with respect to thermal time for shifts between two temperatures in the Arrhenius range 23 °C–37 °C (*n*=253–904 cells). Curves represent the mean, and error bars have been omitted for ease of viewing. G) Single-cell growth rate of *E. coli* MG1655 throughout a temperature upshift from 27 °C to 37 °C in LB (red, *n*=792 cells), casamino acids (green, *n*=270 cells), glucose (blue, *n*=353 cells), glucose+glutamate (orange, *n*=220 cells), or glycerol (purple, *n*=520 cells). Curves show the mean growth rate and the shaded regions represent ±1 SEM. H) Despite the wide variation in absolute growth-rate responses (G), normalized growth rate followed an approximately conserved trajectory versus thermal time across nutrient environments. I) Single-cell growth rates of *E. coli* MG1655 in LB grown to steady state at 37 °C and then subjected to pulses at 25 °C for 10 min (red, *n*=1015 cells), 20 min (orange, *n*=1022 cells), or 30 min (purple, *n*=586 cells). Vertical dotted lines represent the times at which cells were subjected to the downshift to 25 °C for the 10- and 20-min pulses. The upshift from steady-state growth at 25 °C to 37 °C is also shown for comparison (blue, *n*=773 cells). Curves show the mean growth rate and the shaded regions represent ±1 SEM.

To identify the mechanisms underlying the temperature sensitivity of bacterial growth, we used single-cell tracking with a microscopy-compatible temperature controller^32^ to interrogate hypotheses about the molecular architecture underlying temperature sensitivity. For the null model in which growth depends only on a single rate-limiting enzyme, instantaneous growth rate (defined as 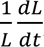, Methods) would adapt immediately to the steady-state value dictated by the current temperature. Hence, the time scale of growth-rate adaptation would match that of a temperature shift. To test whether such a minimal model applies to *E. coli*, we monitored the growth of single cells on LB agarose pads during 10 °C temperature shifts within the Arrhenius range (between 27 °C and 37 °C). A rapid downshift from 37 °C to 27 °C resulted in a rapid decrease in growth rate with a time scale quantitatively similar to that of temperature (Fig. 1B, S3), nearly reaching the steady-state growth rate at 27 °C, and then slowly decelerating toward the new steady state (Fig. 1B).

By contrast, temperature upshifts from 27 °C to 37 °C resulted in a slow and qualitatively distinct response from downshifts, with an initial “spike” that peaked at ∼3 min post-shift and subsequent linear acceleration until reaching the 37 °C steady-state growth rate at ∼40 min post-shift (Fig. 1C), a response time (Methods) substantially longer than the doubling time at 37 °C (∼22 min). To identify factors that affect the response time to temperature upshifts, we shifted *E. coli* from steady-state growth on LB at 27 °C to 30 °C, 33 °C, or 37 °C. In each case, the upshift caused an initial spike and subsequent linear acceleration up to the steady-state growth rate at 37 °C (Fig. 1C,D). However, the time scale of the overall response depended on the final temperature. When cells were shifted to 33 °C, the response time was longer than after an upshift to 37 °C, despite the smaller difference in steady-state growth rate between the initial and final temperature, and the response time for the shift to 30 °C was even longer (∼60 min, Fig. 1D).

To test whether the response is largely determined by the increased growth rate at the elevated final temperature, we normalized time *t* by the doubling time at the final temperature (*τ*_*D*_) to obtain the thermal time *t*/*τ*_*D*_ (Fig. 1E). After normalizing the change in growth rate by the difference in steady-state growth rate at the initial and final temperature, all trajectories collapsed onto a single curve, with response time of 1.6±0.2 doublings at the final temperature (Fig. 1F, Methods). Similar collapse was observed for a broad range of temperature upshifts (Fig. 1F), including to temperatures outside the Arrhenius range (Fig. S4), and across growth media (Fig. 1G,H), suggesting a highly conserved behavior.

Temperature upshift responses were largely unaffected by the presence of oxygen (Fig. S5A,B) or chaperones (Fig. S5C-E). Under nutrient limitation, the enzymes SpoT and RelA produce the alarmone ppGpp, which signals large-scale transcriptional reprogramming known as the stringent response that regulates metabolism^33,34^. During an upshift from 27 °C to 37 °C, the response time of a Δ*spoT* Δ*relA* mutant was longer than the parent strain’s (Fig. S5G), and an upshift from 27 °C to 42 °C caused Δ*spoT* Δ*relA* cells to rapidly decelerate from 0.8 h^−1^ to 0.4 h^−1^ (Fig. S5H), consistent with the transient increase in ppGpp previously observed in these conditions^19,35^. Thus, the stringent response is important for growth at heat-shock temperatures, and its impact on the growth response to upshifts suggests that the response time scale is related to metabolism.

### Downshift pulses reveal a temperature memory

Since adaptation to an upshift requires longer than the doubling time at the final temperature, it is likely that a component(s) limiting for growth must be produced to attain the steady-state growth rate at the higher temperature. In the absence of rapid degradation, after a temperature downshift the level of such a limiting component would represent memory of the growth state at the higher temperature until it is slowly diluted by growth down to its steady-state value at the lower temperature. To evaluate this hypothesis, we monitored MG1655 cells during temperature downshift pulses from 37 °C to 25 °C and back to 37 °C of varying duration. As we hypothesized, after a short (10 min) interval at 25 °C, instantaneous growth rate quickly (within <10 min) recovered back to the 37 °C steady state (Fig. 1I, S6A), accompanied by a larger spike than for cells starting from steady-state growth at 25 °C (Fig. 1I, S6A). For the longest interval at 25 °C (30 min), the response time after the upshift back to 37 °C was still faster (∼25 min, Fig. 1I, S6A) than the steady-state response (∼40 min), indicating that cells did not fully equilibrate to 25 °C 30 min after a downshift from 37 °C. Furthermore, cells shifted back to an intermediate temperature (30°C rather than 37 °C) after 5 min at 23 °C were able to reach the new steady-state growth rate almost immediately (Fig. S6B). Short pulses (5–10 min) at extreme temperatures (<20 °C or >42 °C) from 37 °C generally induced rapid deceleration (Fig. S7), but cells rapidly exited these pulses with large spike responses and re-achieved steady-state growth at 37 °C within 10–20 min (Fig. S7B,C,E). These results suggest that a growth-limiting component is slowly diluted at the lower temperature, as cells can respond quickly to (even extreme) temperature fluctuations.

### The *E. coli* proteome is mostly invariant across temperatures in the Arrhenius range

The prevailing model for growth optimization across nutrient conditions is that growth rate and ribosome concentration are directly coupled^28,36,37^, due to the limitation that ribosomes must produce themselves in addition to all other proteins^26,38^. Within this picture, the rapid response to temperature downshifts and the history dependence of growth rate during downshift pulses might be due to proteins limiting for growth at higher temperatures being overly abundant during the downshift, while the slow response to upshift would be due to the need for proteome reallocation^39^.

To test this hypothesis, we performed untargeted proteomics (LC-MS/MS, Methods) on *E. coli* MG1655 during steady-state growth on three media (LB or minimal media supplemented with glucose or glycerol) at 25 °C, 30 °C, and 37 °C. Of the 4,285 known open reading frames in the *E. coli* MG1655 genome, 1,918 (44.8%) proteins were identified across all samples and biological replicates, similar to the proteome recovery rate of previous studies^40^, accounting for ∼91% of the total proteome estimated from ribosomal profiling (Fig. S8A)^41^ with high replicability (*r*=0.96–1.0) (Fig. S8B,C). We annotated each protein utilizing a modification of the Clusters of Orthologous Groups (COG) database^42^ (Fig. 2A; Methods). As expected, at 37 °C the ribosomal protein fraction was positively correlated with the growth rate in each medium (Fig. 2B, light blue), while energy and metabolism fractions were negatively correlated with growth rate (Fig. 2C, light blue). However, the fraction associated with each functional group was virtually constant across Arrhenius temperatures in each medium (Fig. 2D,E, S8D,E). We only observed significant changes for temperatures outside the Arrhenius range (16 °C, 43 °C) that were consistent with known stress-response pathways (Fig. S8D,F,G)^13^. Thus, ribosome fraction is unlikely to drive growth rate changes across temperatures.

**Figure 2:**
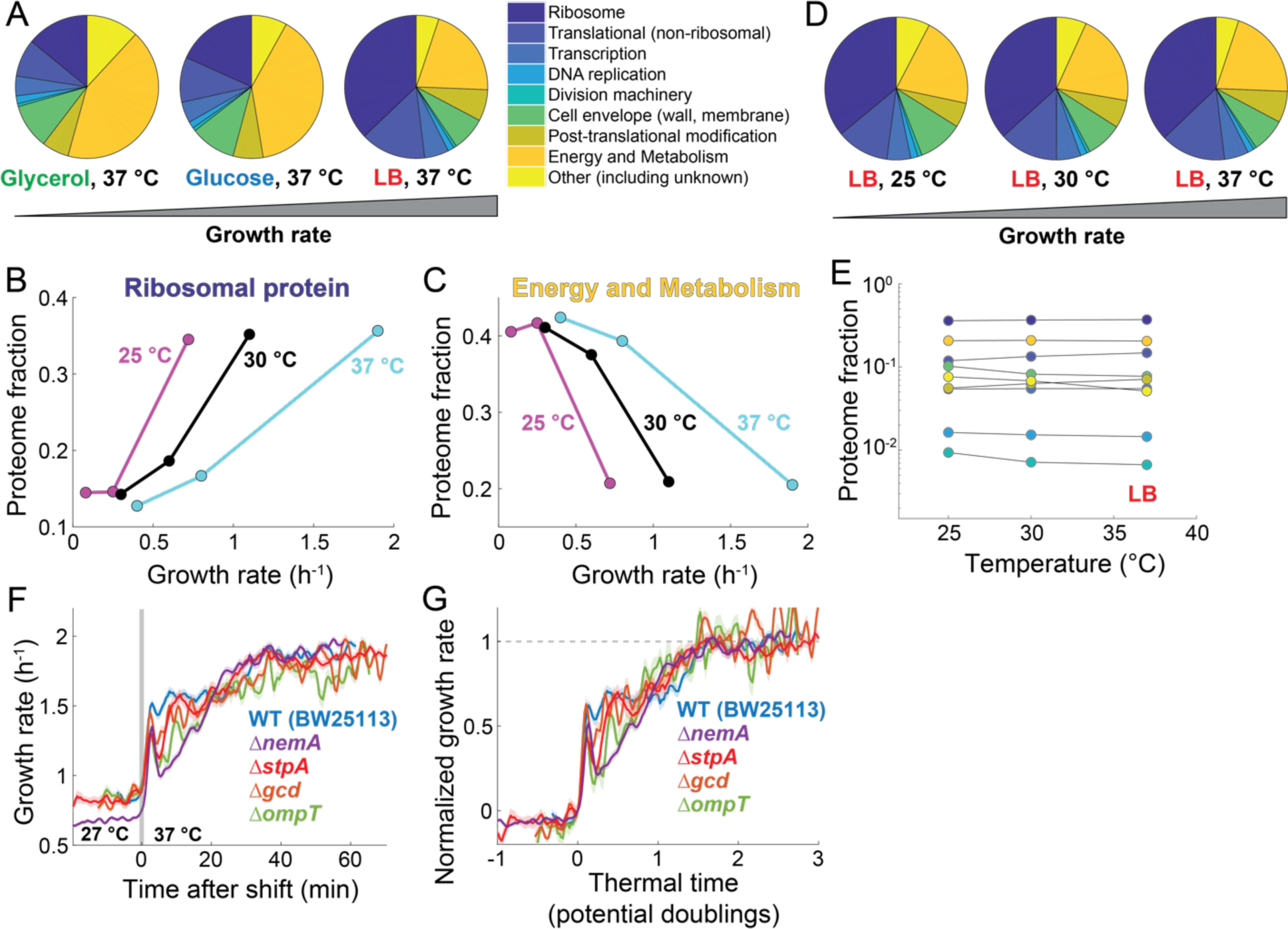
The delayed response to a temperature upshift is not due to proteome rearrangement. A) The *E. coli* proteome at 37 °C varies across media. Functional proteomic sectors (pie chart) were annotated according to the Clusters of Orthologous Groups (COG) classification (Methods). B) Ribosomal protein fraction increased with growth rate across media (M9+glycerol, M9+glucose, LB), independent of growth temperature. C) Energy and metabolism protein fraction decreased with growth rate across media (M9+glycerol, M9+glucose, LB), independent of growth temperature. D) The *E. coli* proteome in LB is largely invariant across temperatures in the Arrhenius range (25 °C–37 °C). E) The proteome fraction accounted for by each functional category was approximately constant in LB across Arrhenius temperatures. Colors correspond to the functions in the legend in (A). F) Temperature-upshift (27 °C to 37 °C) responses of individual deletion mutants of the four most abundant of the proteins whose relative abundance scaled with temperature across all media conditions. (blue: parent strain wild-type BW25113, *n*=710 cells; purple: Δ*nemA*, *n*=1052 cells; red: Δ*stpA*, *n*=588 cells; orange: Δ*gcd*, *n*=814 cells; green: Δ*ompT*, *n*=324 cells). Curves show the mean growth rate and the shaded regions represent ±1 SEM. G) Normalized growth rate followed an approximately conserved trajectory versus thermal time across the mutant growth data in (F).

Most individual protein fractions were constant across temperatures; only 13 proteins exhibited a >2-fold change between 25 °C and 37 °C independent of growth medium (Fig. S9). The most temperature-dependent protein was the DNA-binding StpA^43^, which increased ∼14-fold between 25 °C and 37 °C (Fig. S9A). Nonetheless, Δ*stpA* cells exhibited a similar response to an upshift from 25 °C to 37 °C on LB as wild-type cells (Fig. 2F,G), as did individual knockouts of the three next most temperature-dependent proteins (Fig. 2F,G, S9). Thus, the *E. coli* proteome is largely insensitive to temperature at steady state, and proteins with the largest abundance changes across Arrhenius temperatures do not impact the growth-rate response to temperature upshifts.

### Functional genetic screening did not identify any genes associated with temperature upshifts

While proteomics did not identify large-scale rearrangement between temperatures at steady state (Fig. 2), we interrogated whether some genes determine the non-steady-state dynamics between temperatures (Fig. 1B,C). To test this possibility, we performed a genome-wide, time-resolved screen using a pooled, randomly barcoded transposon mutant library in *E. coli* BW25113^44^, whereby the relative abundance of each mutant was quantified in high throughput during temperature shifts between 25 °C and 37 °C (Fig. 3A, S10A–C,F,G) (Methods). No gene disruptions were consistently identified across biological replicates with a significant fitness defect specific to the temperature upshift (Fig. 3B,C, S10C–E). The only gene of significance from our screen was *uup* (Fig. 3D, S10G), which encodes an ABC-F protein^45^. Δ*uup* cells exhibited slight growth defects at both 27 °C and 37 °C (Fig. 3E), in contrast to other ABC-F mutants (Fig. S10H,I), and the response time after a temperature upshift was substantially longer (∼2.7 doublings versus ∼1.6 for wild type) (Fig. 3F), likely due to defects in ribosome assembly^46^. Taken together, this screen suggests that none of the nonessential genes are responsible for the delayed growth-rate response to temperature upshifts.

**Figure 3:**
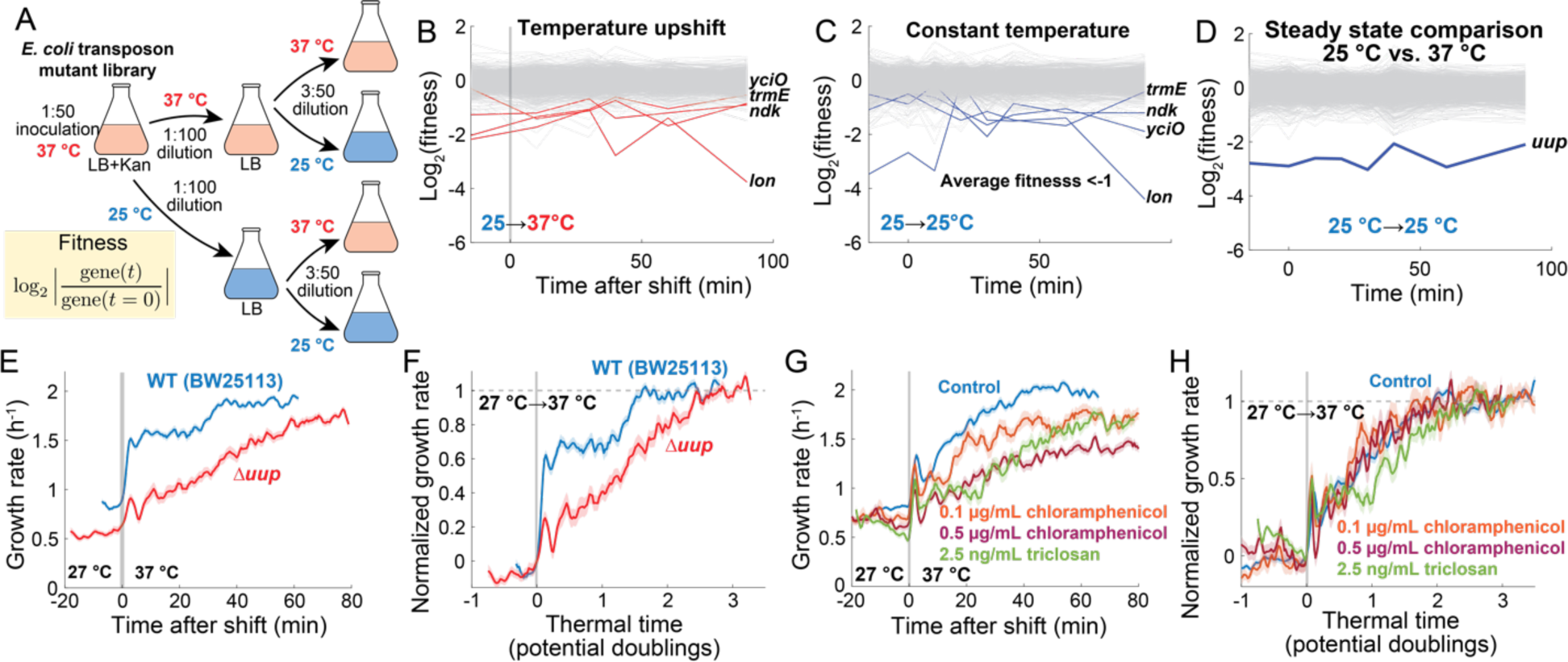
Response time is unaffected by almost all genetic and chemical perturbations. A) Schematic of temperature-shift experiments screening a pooled library of *E. coli* BW25113 randomly barcoded transposon mutants (Methods). The library was initially grown in a 50-mL volume at 37 °C, then diluted and grown to steady state at the initial target temperature (25 °C or 37 °C) before shifting to 25 °C or 37 °C at time *t*=0. The optical density was monitored at each sampling time point (∼10 min intervals). The fitness of each mutant is defined as the log_2_(relative abundance compared to *t*=0) (Methods). This experiment was carried out twice. B) Trajectories of mutant fitness during a shift from 25 °C to 37 °C (gray vertical bar denotes timing of the shift). Mutants with fitness<-1 when averaged over time points are highlighted in red. C) Trajectories of mutant fitness during steady-state growth at 25 °C. Mutants with fitness<-1 when averaged over time points are highlighted in blue. (B) and (C) are from the same experiment, and outlier mutants in the upshift (B) were also outliers in the control (C). D) Trajectories of log_2_(relative abundance) during steady-state growth at 25 °C compared with at 37 °C. A single gene, *uup*, had fitness<-1 when averaged over time points. E) Single-cell growth rate of Δ*uup* (red, *n*=609 cells) and its parent BW25113 (blue, *n*=734 cells) throughout a temperature shift from 27 °C to 37 °C. Curves show the mean growth rate and shaded regions represent ±1 SEM. F) Normalized growth rate versus thermal time for each trajectory in (E) shows that Δ*uup* cells respond more slowly to an upshift. G) Single-cell growth rates of *E. coli* MG1655 in LB treated with 0.1 μg/mL chloramphenicol (orange, *n*=264 cells), 0.5 μg/mL chloramphenicol (dark red, *n*=382 cells) or 2.5 ng/mL triclosan (green, *n*=266 cells) throughout a shift from 27 °C to 37 °C. The untreated control is also shown for comparison (blue, *n*=792 cells). Curves show the mean growth rate and shaded regions represent ±1 SEM. H) Normalized growth rate followed a similar trajectory versus thermal time as the control for the chloramphenicol treatment data in (G), while triclosan slightly delayed the growth-rate response.

To investigate the role of essential genes during temperature upshifts, we treated *E. coli* cells with a variety of antibiotics at sub-minimum inhibitory concentration (MIC) levels^47^ and subjected them to temperature upshifts from 27 °C to 37 °C on LB (Fig. S11). Response times remained ∼1.5 doublings for treatment with antibiotics that target the ribosome (Fig. 3G,H, S11A,B), DNA replication, and transcription, despite large impacts on growth rate in some cases (Fig. S11C,D). Only at high concentrations of fusidic acid (50 μg/mL), which disrupts translational translocation and ribosome disassembly^48^, did we observe a significant decrease in response times (∼0.2 doublings; Fig. S11F,G). We observed similar effects in knockouts of genes responsible for tRNA modification (*tusA*, *tusB*)^49^ (Fig S11F,G), suggesting that severely impaired translational capacity under high amino acid availability causes a mismatch, allowing for fast response times.

Membrane fluidity is exquisitely regulated in *E. coli* across temperatures: the fraction of unsaturated fatty acids decreases with increasing temperature to maintain viscosity^24,25^. Triclosan, which targets membrane synthesis, increased the response time to ∼2.5 doublings (Fig. 3G,H), suggesting that the delay in growth rate might be due to the need to alter membrane composition. However, disruptions to the regulation of *fabB*, which encodes the major β-ketoacyl-[acyl carrier protein] synthase responsible for elongating unsaturated fatty acids^50,51^, had little effect on upshift response times, despite significant changes to membrane fluidity (Fig. 4A–D) (Methods). While an increase in membrane fluidity conferred a growth advantage at 37 °C (Fig. 4F), we found that increases in membrane fluidity negatively affected cell growth and survival at temperatures >42 °C (Fig. 4G,H). Thus, while membrane fluidity does not determine the time scale of upshift responses, its regulation is critical for cell integrity at high temperatures.

**Figure 4:**
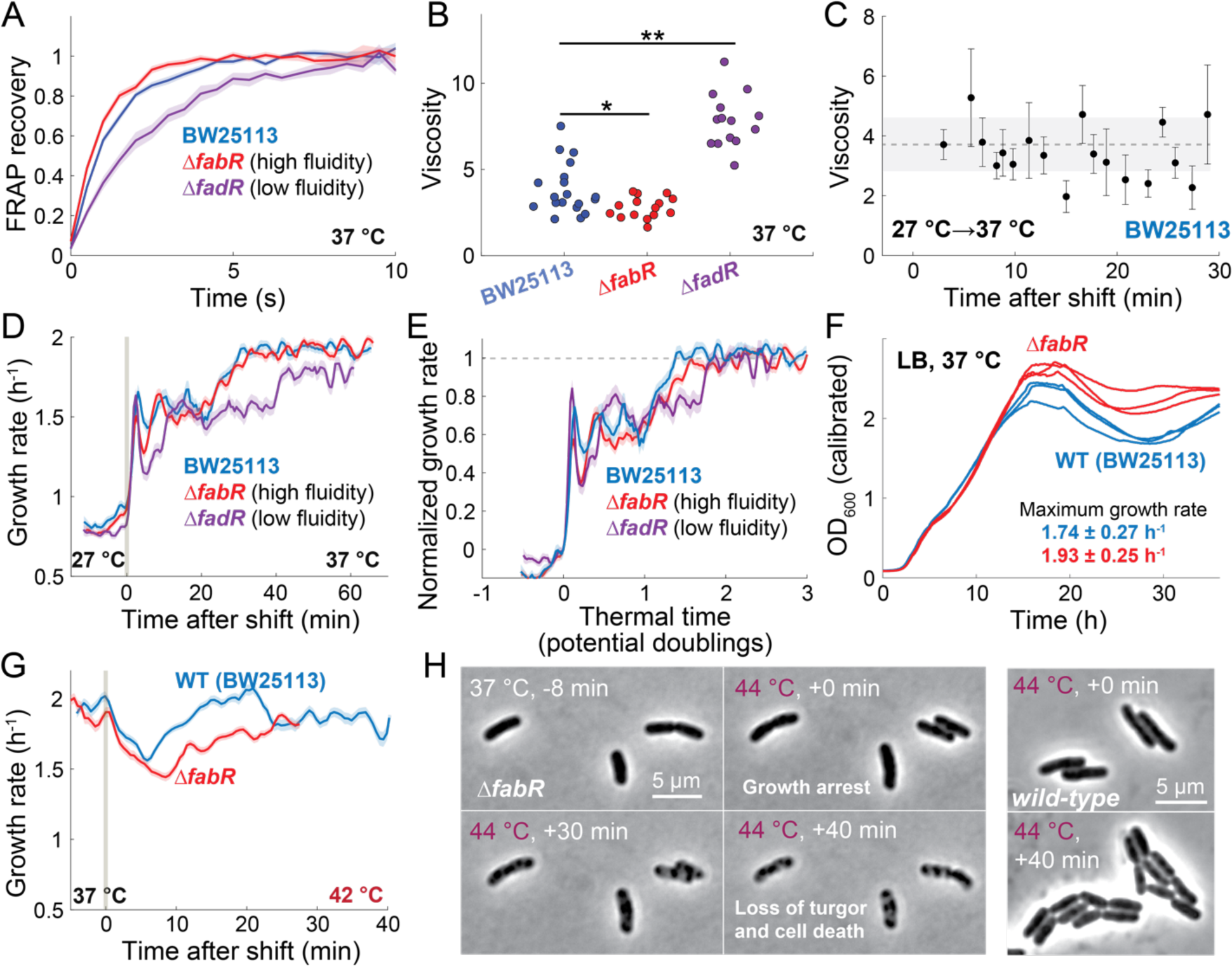
Changes in membrane fluidity do not alter the response to temperature upshift but increase lysis at high temperatures. A) FRAP recovery dynamics of the membrane dye MitoTracker in the parent strain BW25113 (blue, *n=*19 cells), the high-fluidity mutant Δ*fabR* (red, *n*=15 cells), and the low-fluidity mutant Δ*fadR* (purple, *n*=14 cells) grown on LB (Methods). Curves show the mean recovery and shaded regions represent ±1 SEM. B) Membrane viscosity (Methods) was significantly lower and higher in Δ*fabR* and Δ*fadR*, respectively, compared with the parent. Significance was determined using a two-sample t-test; *: *p*<0.05, **: *p<*0.005. C) Membrane viscosity of *E. coli* BW25113 cells after an upshift at *t*=0 from 27 °C to 37 °C in LB (Methods). Points are estimates from a best fit (Methods) and error bars represent 1 standard error. Dotted line is the mean viscosity from steady-state measurements (B), and the shaded region represents ±1 standard deviation. D) Growth rate responses of *E. coli* BW25113 (blue, *n*=686 cells), Δ*fabR* (red, *n*=924 cells), and Δ*fadR* (purple, *n*=409 cells) to a temperature upshift from 27 °C to 37 °C in LB. Curves show mean growth rate and shaded regions represent ±1 SEM. E) Normalized growth rate followed a similar trajectory versus thermal time among the mutants and parent for the data in (D). F) Growth curves of wild-type (BW25113) (blue) and Δ*fabR* (red) cells grown in LB at 37 °C (*n*=3 replicates). Optical density (OD) was corrected for non-linearity at high OD values (Methods). Maximum growth rates are the mean across replicates and the error is ±1 standard deviation. G) Growth rate response to a temperature upshift from 37 °C to 42 °C in *E. coli* BW25113 (blue, *n*=1011 cells) and Δ*fabR* (red, *n*=554 cells). Both strains exhibited an initial decrease in growth rate followed by recovery to the steady-state growth rate at 37 °C, but recovery was more delayed for the high-fluidity Δ*fabR* mutant. Curves show mean growth rate and shaded regions represent ±1 SEM. H) (Left) Representative images of Δ*fabR* cells throughout a temperature upshift from 37 °C to 44 °C. At 37 °C, morphology and growth were wild-type-like (top left). Growth halted immediately after the shift to 44 °C (top right), with loss of turgor and cell death occurring within 30–40 min after the shift (bottom left, right). (Right) Representative images of wild-type MG1655 cells before and after an upshift from 37 °C to 44 °C. Cells maintained growth and shape.

### A temperature-sensitive enzyme network (TSEN) model captures temperature-shift response dynamics

Our findings indicate that the normalized response time to temperature upshifts is largely conserved (Fig. 1F,H, 2G, 3H, 4E, S5B,E, S10I, S11B,D). Our proteomic, genetic, and chemical screens largely failed to uncover a key molecular regulator of the response, leaving it unclear how the response behavior arises and what factors set the time scale. To interrogate the mechanisms underlying these phenomena, we developed an autocatalytic network model of growth, since such models have a long history in enzyme kinetics^15^.

To model growth, we incorporated three general classes of reactions: (1) import, (2) metabolite production, and (3) volume expansion (Fig. 5A). Each reaction obeys Michaelis-Menten kinetics, such that metabolite *c_i_* is consumed by enzyme *e_i_* to produce the next metabolite *c_i_*_+1_ in the network:

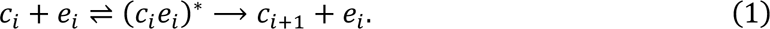

**Figure 5:**
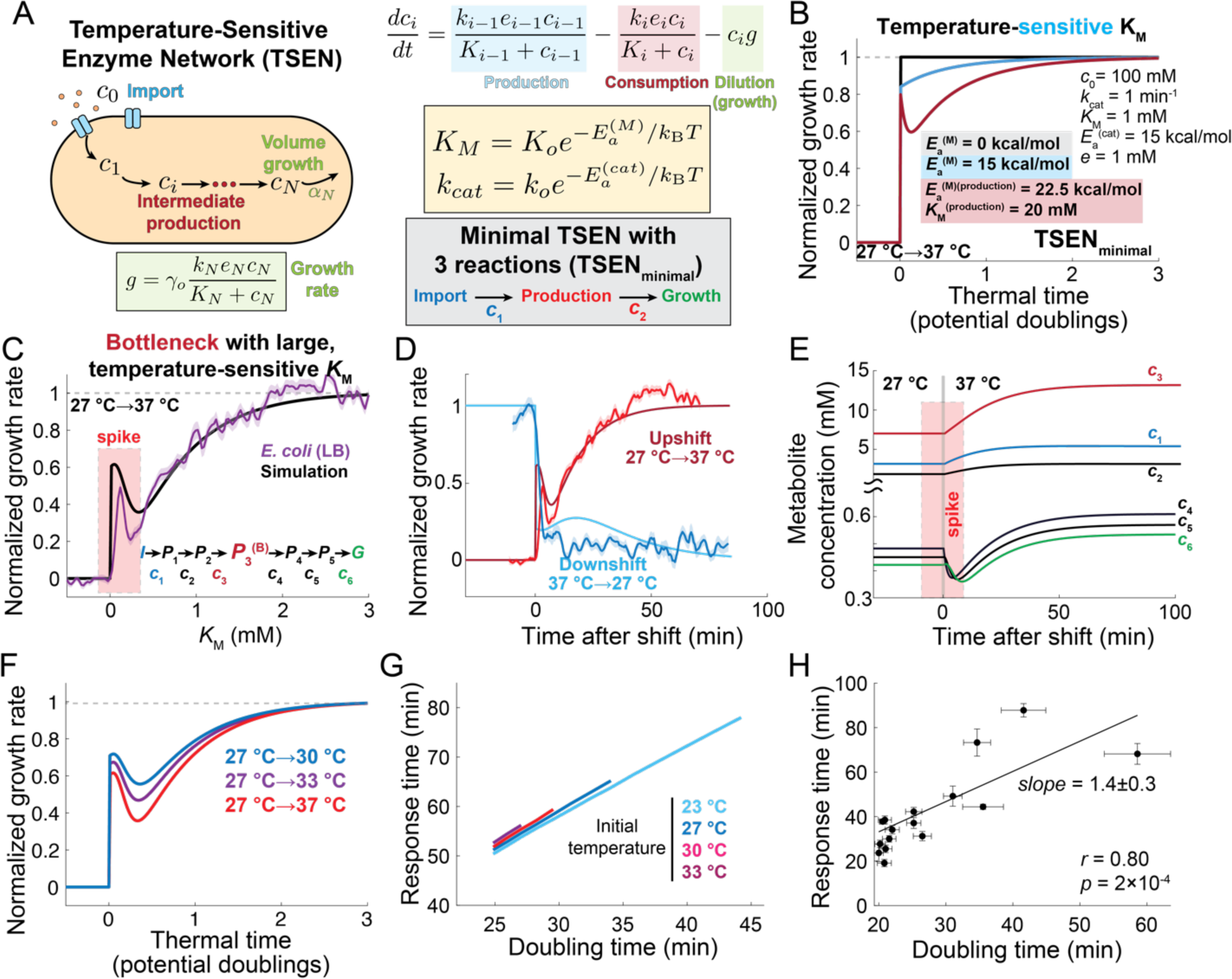
A temperature-sensitive enzyme network (TSEN) model recapitulates temperature-shift behaviors through metabolome rearrangement. A) (Upper left) In the TSEN model, the import reaction (blue), production reactions (dark red), and volume growth reaction (green) are chained Michaelis-Menten reactions. (Upper right) The rate equation for each intermediate metabolite (*c_i_*) involves enzymatic production from *c_i_*_-1_ by enzyme *e_i_*_-1_ (blue), enzymatic consumption by enzyme *e_i_* (red), and dilution by volume growth *g* (green). The final intermediate *c_N_* is translated into volume growth with an efficiency factor *γ*_0_(green box). Each kinetic parameter is assumed to be temperature sensitive according to an Arrhenius equation with activation energy *E_a_* (yellow box). A minimal TSEN model (TSEN_minimal_) has a single intermediate production reaction (red), such that the network has only 2 intermediate metabolites (*c*_1_, *c*_2_; gray box). B) Normalized growth rate versus thermal time from simulations of a minimal TSEN model with (blue) and without (black) a temperature-sensitive Michaelis-Menten constant (K_M_) throughout a temperature shift from 27 °C to 37 °C. All other model parameters are identical and are defined in the panel inset. The TSEN with a temperature-sensitive K_M_ results in a non-zero response time (blue). A TSEN model with a bottleneck (dark red, *K_M_* = 20 mM, *E_α_* = 22.5 kcal/mol) produces an initial spike. C) A TSEN model with a single bottleneck (red, *P*_3_) embedded within a 7-reaction chain produces a quantitatively similar response to an upshift from 27 °C to 37 °C as *E. coli* MG1655 cells on LB (purple, *n*=792 cells; shaded region represents ±1 SEM). All other reactions have parameters identical to the minimal model without a bottleneck (blue) in (B). D) Normalized growth rate in simulations of the bottlenecked TSEN model in (C) throughout an upshift (dark red) or downshift (light blue) between 27 °C and 37 °C. In contrast to the spike and slow response to the upshift, the downshift results in immediate deceleration to a growth rate close to the steady-state value at 27 °C, followed by slow deceleration to new steady-state value. Overlaid are the normalized growth rates of *E. coli* MG1655 subjected to an upshift (red, 27 °C to 37 °C, *n*=792 cells) or downshift (blue, 37 °C to 27 °C, *n*=253 cells). Shaded error bars represent ±1 SEM. E) The TSEN model in (C) predicts that metabolites upstream of the bottleneck (*c*_1,2,3_) will have lower concentrations than those after the bottleneck (*c*_4,5,6_) and increase slowly on the time scale of growth after an upshift. Post-bottleneck metabolites undergo transient decreases immediately after the temperature upshift over a time interval corresponding to the spike (red box). After this initial period, all metabolites increase to new values corresponding to the steady state at the higher temperature. F) The TSEN model in (C) predicts highly similar responses of normalized growth rate with thermal time for upshifts from 27 °C to 30 °C (blue), 33 °C (purple), or 37 °C (red). G) Over a broad range of final temperatures, the absolute upshift response time (time to reach 98% of the difference between steady-state growth rates) has linear scaling with the doubling time at the final temperature. Each line represents simulations from a given initial temperature to temperatures up to 37 °C. H) Experimental estimates of absolute response time follow a linear scaling with doubling time at the final temperature, similar to simulations in (G). Shown are data for upshifts involving initial and final temperatures ranging between 23 °C and 37 °C (data from Fig. 1F–H).

In a linear network (no branching), the dynamics of each intermediate metabolite are dictated by

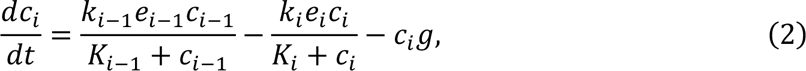

where the first two terms on the right-hand side reflect enzymatic production and consumption of *c_i_* and the last term accounts for dilution via growth at the rate *g* (SI). *e_i_* is the concentration of the *i*^th^ enzyme, *k_i_* is its catalytic rate, and *K_i_* is its Michaelis-Menten constant. Importantly, we consider the possibility that both *k_i_* and *K_i_* are temperature dependent with Arrhenius behavior (Fig. 5A, SI), consistent with experimental measurements of several enzymes^52–54^. After the final step of a reaction network of size N, an enzyme *e*_N_ consumes *c*_N_ to expand cell volume at a rate *g* = 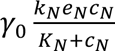, where *γ*_0_ is an efficiency factor assumed to be constant in a given environment, reflecting the conversion of *c*_N_ to a structural component of the cell envelope (SI). From physical considerations of glucose uptake and growth rate constraints (SI), we estimated that *γ*_0_ is ∼0.03 mM^−1^. We call these equations a temperature-sensitive enzyme network (TSEN) model.

Since the proteome is largely maintained across temperatures (Fig. 2E, S8D), we assumed that the concentration of each enzyme remains constant during any temperature shift. In a simple version of our model with two intermediate metabolites (minimal TSEN), we considered a single intermediate reaction between import and growth and set each kinetic parameter to be identical across reactions (Fig. 5A). We also assumed a high (saturating) concentration of the external substrate. At a given temperature, the system reaches a steady state with stable concentrations of the intermediate metabolite.

We simulated the model to reach a steady state at 27 °C and then shifted the temperature to 37 °C. The system exhibited a non-zero response time to reach the steady-state growth rate at 37 °C (Fig. 5B). To increase growth rate by 95% of the difference between 27 °C and 37 °C required ∼2 doublings (Fig. 5B), similar to our experimental observations (Fig. 1F,H). Variations in kinetic parameters largely maintained response times of 1–2 doublings (Fig. S12A,B), as long as the *E_α_* for the catalytic rate of the import reaction was above that of the other catalytic rates (production, growth) (Fig. S12B). However, if the activation energy for all Michaelis-Menten constants is 0, then the growth rate response is also immediate (Fig. 5B), highlighting the importance of temperature sensitivity of *K*_&_. Furthermore, the version of the minimal TSEN without an intermediate reaction (“production-less” TSEN, SI) is analytically tractable and predicts an upshift response time between 1.1 and 2.3 doublings for a growth-rate cut-off between 95% and 99% of the difference between steady-state growth rates (Fig. S12C–F), in agreement with our measurements (∼1.5 doublings, Fig. 1F,H), suggesting that the precise definition of the response time scale is not critical for producing physiological timescales. Additionally, the minimal TSEN produces an Arrhenius-like growth rate (Fig. S12G), in agreement with observations^10,12^ (Fig. 1A, S1, S2).

For a reaction network with N intermediates, the analytical solution of steady-state growth rate (SI) is

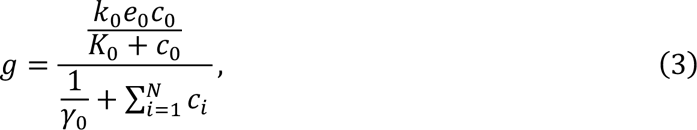

which is set by the import rate 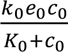, growth efficiency factor *γ*_0_, and the total intracellular metabolite concentration 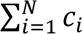. Since import changes instantaneously with temperature, the response time scale is associated with rearrangement of the metabolome. Eq. 3 also reflects that growth comes at the cost of storing metabolic intermediates *c_i_*, and holds regardless of branching within the network (SI). This model recovers the well-known Monod Equation as long as import follows Michaelis-Menten kinetics^55^ (SI), which has been shown for a variety of sugar transporters in E. coli^56–58^.

In *E. coli*, the majority of major metabolites are present at concentrations that saturate their respective enzymes^59^. However, many central-carbon reactions operate near their *K_M_*^59^, suggesting that bottlenecks are highly likely. Moreover, enzymes that catalyze product formation tend to have higher activation energy than transporters^60^. Within a simple network involving import, production, and growth, introducing a bottleneck into the production reaction characterized by a large and temperature-sensitive *K_M_* (*K_M_*(*T* = 37 °C) = 20 mM, *E_a_*=22.5 kcal/mol) produced strikingly physiological behavior: simulations predicted an initial spike similar to our experimental observations and a response time of ∼2 doublings (Fig. 5B). Moreover, when the bottleneck was within a pathway involving multiple production reactions, the spike broadened, and the acceleration dynamics were quantitatively similar to our experimental measurements (Fig. 5C). This TSEN model readily predicts asymmetric behavior between upshifts and downshifts (Fig. 5D); such asymmetry is expected from temperature-dependent Michaelis-Menten kinetics^12^. Additionally, after the initial large decrease in growth rate upon a downshift, the multiple-reaction model predicts a slow deceleration toward the slower steady-state growth rate (Fig. 5D, dark blue), similar to our experimental observations (Fig. 1C, S2). Thus, an initial spike, a physiological response time, and asymmetric behavior emerge from only a few simple assumptions about the network.

### Spike and response time are dictated by metabolome rearrangement

In the bottlenecked TSEN model, metabolites after the bottleneck undergo a transient decrease in concentration after a temperature upshift (Fig. 5E) that agrees with the time scale of the spike response (Fig. 5D), suggesting that spikes are caused by rapid consumption of post-bottleneck metabolites. Meanwhile, metabolites before the bottleneck gradually build up, as do the post-bottleneck metabolites after the spike, until metabolome rearrangement has stabilized (Fig. 5E). As a result, final growth rate (which scales with temperature) dictates the thermally limited response time necessary to rearrange the metabolome. Simulations of our model captured the near invariance of the normalized response time across initial and final temperatures (Fig. 5F) as represented by linear scaling of absolute response time with doubling time at the higher temperature (Fig. 5G). This scaling was consistent with our experimental measurements (Fig. 2E), which exhibited a slope of 1.4±0.3 (Fig. 5H), consistent with the range of normalized response times measured across temperatures and growth media (Fig. 1F,H).

### TSEN model predicts spike dependence on nutrient type

Early work by Monod and others demonstrated that *E. coli* growth rate depends on glucose concentration in a manner remarkably similar to Michaelis-Menten enzyme kinetics with a half-saturation constant *K_M_*∼22 μM^55^. This relationship is referred to as the Monod Equation,

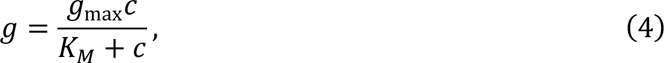

where *c* is the concentration of the substrate (e.g., glucose) and *g*_max_ is the growth rate when *c* ≫ *K_M_*. This equation is highly similar to the steady-state growth prediction from the TSEN model (Eq. 3), which predicts that *g*_max_ and *K_M_* are largely determined by import, which in turn likely depends on the specific substrate. To examine how these temperature-dependent parameters vary with growth substrate, we measured the growth rate *g*_max_ and *K_M_* at various temperatures across a variety of media, including simple sugars, casamino acids, and rich medium (LB) (Fig. S13A–C; Methods). Our measurements showed that growth on simple sugars produces Monod-like behavior with large *E_α_* for *g*_max_ (12–25 kcal/mol) and small, temperature-insensitive *K_M_* (0.04–0.3 mM) (Fig. S13C–F). Conversely, amino acid growth was characterized by smaller, concentration-dependent *E_α_* for *g*_max_ (3–15 kcal/mol) with large, temperature-sensitive *K_M_* (1–9 mM) (Fig. S13C–G).

Despite the higher temperature sensitivity (*E_a_*) of steady-state growth rate in glucose compared with casamino acids or LB (Fig. S13), the normalized response time for an upshift was highly similar across these nutrients (Fig. 1H), as predicted by our TSEN model, wherein increases in the activation energy of the import reaction have little effect on response times (Fig. S12B). However, the spike was much smaller or nonexistent in glucose (Fig. 1H). Motivated by this decoupling of response time and spike height, we probed the behavior of our TSEN model when changing the activation energy of the catalytic rate of the import reaction to mimic the differences between LB and glucose. The TSEN model predicts that spike height should decrease sharply when the activation energy of import is >15 kcal/mol (Fig. 6A), while the response time increases by only ∼0.1 doubling (Fig. 6B). Thus, the model predicts that activation energy differences between nutrient types (Fig. S13F,G) are sufficient to explain differences in the initial spike behavior observed across growth media (Fig. 2G,H).

**Figure 6:**
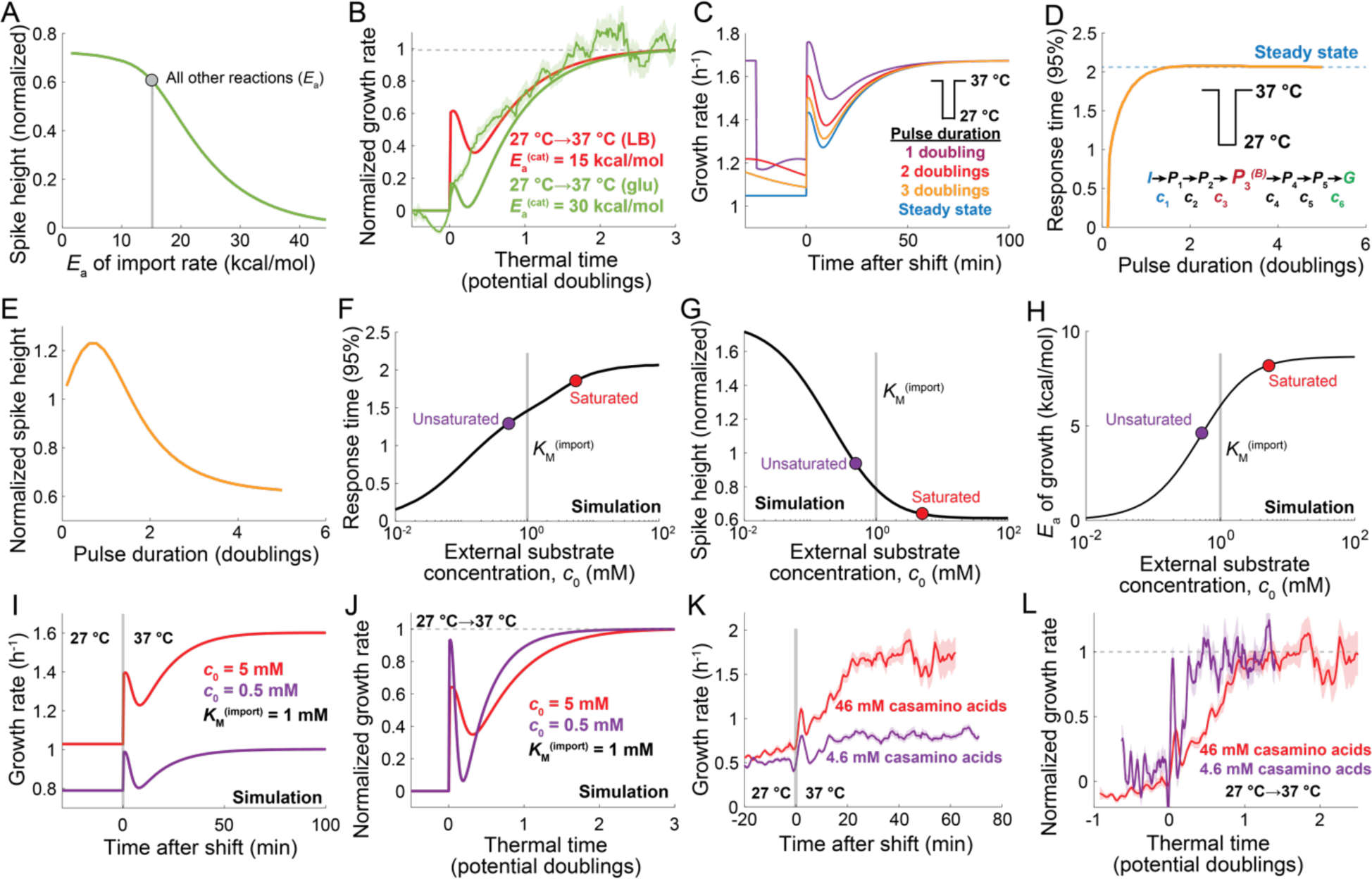
TSEN model predicts changes to upshift response during growth on simple sugars, after a downshift pulse, and at low nutrient concentration. A) The bottlenecked TSEN model in Fig. 5C predicts that spike height after a temperature upshift will decrease with the activation energy of import. The activation energies of all other reactions were set to 15 kcal/mol (gray circle). B) Increasing the activation energy of import to *E_α_* = 30 kcal/mol based on our experimental measurements of the activation energy of growth in glucose (Fig. S12G) dramatically reduced the spike height (green) compared with our default *E_α_* = 15 kcal/mol (red) without affecting the response time, similar to our experimental measurements of the normalized growth-rate response on glucose (green data, same as in Fig. 1H). C) The bottlenecked TSEN model predicts that short downshift pulses from 37 °C to 27 °C (purple, red, orange) will cause larger spikes and faster recovery to the steady-state growth rate at 37 °C than an upshift from steady state at 27 °C to 37 °C (blue), similar to our experimental observations (Fig. 1I). D,E) The bottlenecked TSEN model in Fig. 5C predicts that the response time decreases for downshift pulses of duration <1 doubling at the higher temperature (D) and that the spike height remains high for pulses of duration multiple doublings at the higher temperature (E). F,G,H) The bottlenecked TSEN model in Fig. 5C predicts that response time decreases (F), spike height increases (G), and growth activation energy decreases (H) when nutrient concentration is reduced from above (saturated, red circle) to below (unsaturated, purple circle) the K_M_ for import (gray vertical bar). I) Simulations of the bottlenecked TSEN model in Fig. 5C throughout an upshift from 27 °C to 37 °C for high (*c*_0_=5 mM, red) or low (*c*_0_=0.5 mM, purple) external substrate concentration (K_M_(import) = 1 mM). The absolute spike height is similar in magnitude at both concentrations, but the steady-state growth rate at 37 °C is much lower for lower external substrate concentration (purple) and is reached more quickly after the upshift than for higher concentration (red). J) The normalized growth rate calculated from simulations in (H) has a larger spike and a faster normalized response (<1 doubling) for low external substrate concentration (purple) compared with the TSEN model at saturation (red). K,L) Growth rate throughout an upshift from 27 °C to 37 °C of *E. coli* MG1655 grown on MOPS+46 mM casamino acids (red, *n*=116 cells) or 4.6 mM casamino acids (purple, *n*=261 cells) exhibits similar dynamics in both absolute (J) and normalized (K) terms as the simulations in (H,I). Curves are mean growth rate and shaded regions represent ±1 SEM. The response time at low casamino acid concentration is much shorter (purple, ∼0.25 doublings) compared to at saturation (red, ∼1.4 doublings) and is accompanied by a larger relative spike height.

### TSEN model predicts a metabolically encoded temperature memory

While growth rate rapidly decreases after a downshift in our model (Fig. 5D), the concentration of metabolites requires dilution to equilibrate, which takes place on the time scale of growth. Thus, during a transient downshift pulse, the cell gradually transitions from the high temperature to the low temperature metabolic state, despite rapid growth deceleration. Indeed, simulations of the bottle-necked TSEN model (Fig. 5C) during a downshift pulse from 37 °C to 27 °C back to 37 °C, with variable duration at 27 °C, resulted in faster upshift responses than from the steady state at 27 °C (Fig. 6C). More than a doubling at 27 °C was required for recovery of the upshift response time (Fig. 6D) and the spike height remained higher than for an upshift from steady state for pulses of multiple doublings (Fig. 6E), again collectively consistent with our experimental data (Fig. 1I, S7). These findings highlight the ability of our model to reproduce nearly all observed temperature-shift responses with reasonable quantitative agreement, underscoring the importance of the temperature sensitivity of import and metabolome rearrangement in upshift responses.

### TSEN model successfully predicts temperature upshift response at low substrate concentration

Although we have thus far focused on experimental conditions in which the external substrate concentration is saturating, nutrient concentrations are low in many environments. Since a key feature of the ability of our model to successfully predict upshift responses is the temperature sensitivity of Michaelis-Menten constants, we were motivated to examine the low-nutrient regime. To examine this regime, we simulated our model with external concentrations below the *K_M_* for import and considered a temperature upshift from 27 °C to 37 °C. At *c*_0_ < *K*_0_, the relative spike height increased substantially, and the normalized response time decreased with decreasing concentration (Fig. 6F–J). This decrease in response time occurred because the increase in import rate after a temperature upshift is smaller when *c*_0_ ≤ *K*_0_ compared with *c*_0_ ≥ *K*_0_^12^, despite an identical increase in catalytic rates that consume the imported metabolite. The activation energy for growth was also predicted to decrease with decreasing nutrient concentration (Fig. 6H), in agreement with our experimental measurements of growth on amino acids (Fig. S13G).

To test these predictions, we shifted MG1655 cells from 27 °C to 37 °C during growth on casamino acids (Methods). With 4.6 mM CAA, a concentration near the *K_M_* (Fig. S13C), growth rate increased from the steady state value at 27 °C (∼0.5 h^−1^) to that at 37 °C (∼0.8 h^−1^) after only 25 min, corresponding to 0.25 doubling times at 37 °C (Fig. 6K,L). Moreover, the spike peaked at a growth rate close to that of the 37 °C steady state (Fig. 6K). These dynamics were in reasonable agreement with the predictions of our model (Fig. 6I,J), indicating that the response to temperature shift can be modulated by nutrient concentration, owing to the properties of the import *K_M_*.

### Temperature shifts in fission yeast produce a distinct response from bacteria that is captured by the TSEN model

Diverse bacterial species, including *E. coli* from host organisms with distinct temperature-dependent evolutionary histories (turtle, seagull, human) and the soil-dwelling gram-positive *Bacillus subtilis*, exhibited similar responses to upshifts as *E. coli* MG1655 (Fig. 7A,B, S14), suggesting a general behavior across the bacterial kingdom. We sought to determine if single-celled eukaryotes would respond to temperature shifts in a similar manner. The rod-shaped fission yeast *Schizosaccharomyces pombe* grows optimally at 32 °C and possesses an Arrhenius range between 22 °C and 32 °C, with *E_a_*∼8 kcal/mol in rich media^61^. During an upshift on the rich medium YE5S from 22 °C to 32 °C, cells exhibited a large growth-rate spike from 0.17 h^−1^ to 0.44 h^−1^, overshooting the steady-state growth rate at 32 °C (Fig. 7C). This spike was immediately followed by a deceleration to 0.02 h^−1^, followed by rapid acceleration to the steady-state growth rate at 32 °C (∼0.4 h^−1^) (Fig. 7C). The entire response dynamics lasted ∼100 min, corresponding to a normalized response time of ∼1.1 doublings (Fig. 7D). Thus, despite decelerating to near growth halting, *S. pombe* cells were able to reach the steady-state growth rate at the higher temperature more quickly than any bacteria tested at saturating nutrient concentration.

**Figure 7:**
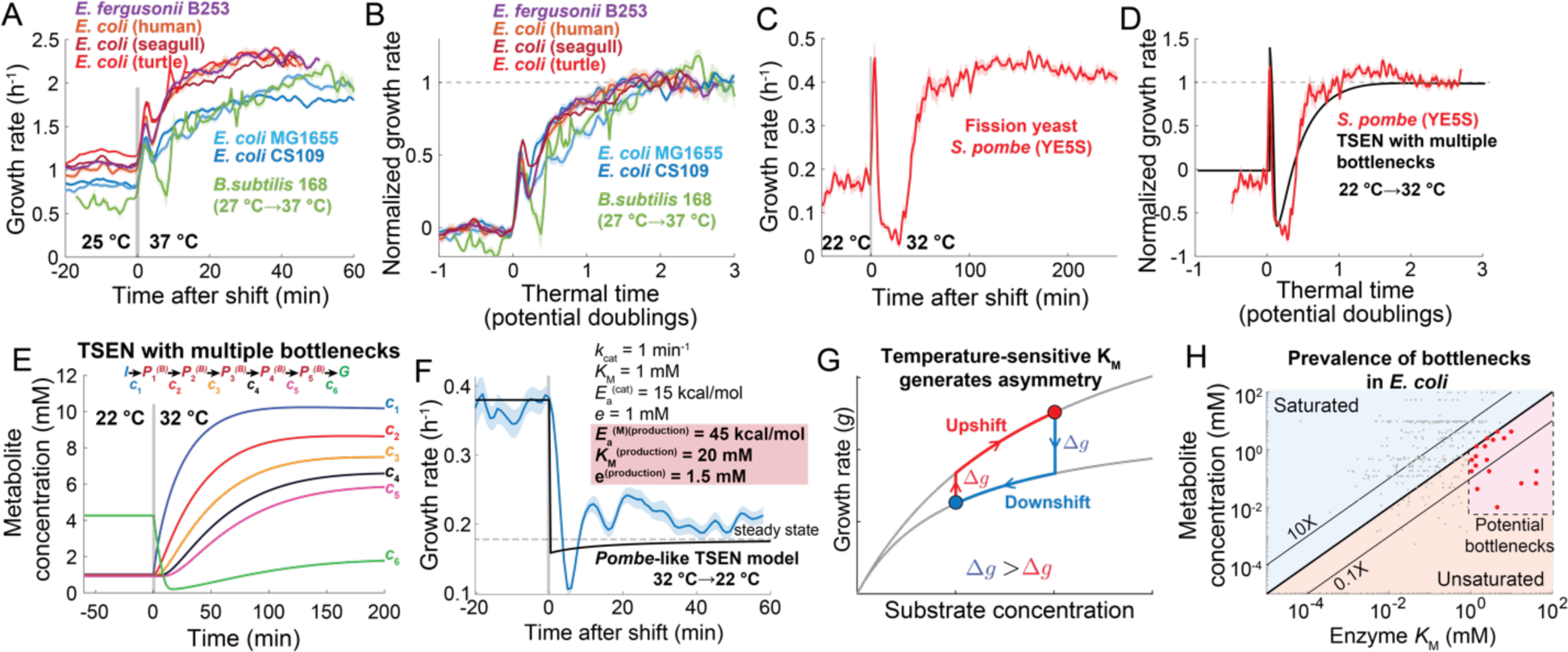
Temperature upshift response features are generally conserved across microbes and can be captured by the TSEN model. A) Growth-rate response to a temperature upshift on LB of laboratory-evolved *E. coli* (light and dark blue, *n*=773–1279 cells) and natural isolates from various hosts (orange to red, *n*=389–547 cells), *Escherichia fergusonii* (purple, *n*=997 cells), and *Bacillus subtilis* (green, *n*=236 cells). All upshifts were from 25 °C to 37 °C, except for *B. subtilis* (27 °C to 37 °C). Curves are mean growth rate and shaded regions represent ±1 SEM. B) Normalized growth rate followed a similar trajectory versus thermal time across all strains/species for the data in (A). C) A temperature upshift from 22 °C to 32 °C on the rich growth medium YE5S caused the growth rate of the fission yeast *Schizosaccharomyces pombe* to initially spike to close to the steady-state value at 32 °C, then decelerate to below the steady-state value at 22 °C, then accelerate back to the new steady-state value within less than a doubling (∼100 min) (red, *n*=333 cells). Curves are mean growth rate and shaded regions represent ±1 SEM. D) A TSEN model with 5 intermediate reactions, each with a large (20 mM) and highly temperature-sensitive K_M_ (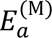 = 22.5 kcal/mol) produces similar normalized growth rate dynamics with thermal time (black) as the *S. pombe* data in (C) (red), characterized by a large spike, deceleration to <0, and fast recovery within <1 doubling. E) The TSEN model in (D), composed entirely of bottlenecks, predicts that a temperature upshift results in a rapid, large decrease in the final intermediate metabolite concentration (*c*_6_), followed by a slower increase to the new steady-state, which is lower than at lower temperature. All other metabolite concentrations (*c*_1-5_) increase monotonically toward their steady-state levels at rates that depend inversely on their network depth (i.e., *c*_5_ increases more slowly than *c*_1_). F) An extended TSEN model with five intermediate reactions, each with a large (20 mM) and highly temperature-sensitive (E_a_=22.5 kcal/mol) K_M_, predicts a temporary undershoot (black) upon a downshift to growth rates below the steady state value at the lower temperature. The fission yeast *Schizosaccharomyces pombe* exhibited an undershoot during a downshift from 32 °C to 32 °C on rich medium (YE5S) (blue, *n*=54 cells) similar to model predictions. Experimental data are the mean±1 SEM (shaded region) at each time point. G) Asymmetry in growth-rate response to temperature up- and downshifts is caused by the temperature sensitivity of K_M_. If substrate concentration is near the K_M_ and 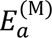 > 0, an upshift causes a small initial increase in growth rate, followed by a slow increase in substrate concentration to the new K_M_ via production (red). Conversely, a downshift causes a comparatively larger initial decrease in growth rate, followed by a slow decrease in substrate concentration to the new K_M_ via dilution (blue). H) Intracellular metabolite concentrations are generally much higher than the K_M_ of their production enzyme for *E. coli* grown on glucose (data from Ref. ^81^). However, some metabolites have concentrations near or below the K_M_ of their production, and potential bottlenecks (purple region, red points) are those with K_M_>1 mM and substrate concentration *c*<K_M_.

We wondered whether these differences in behavior were consistent with our TSEN model. By systematically varying parameters, we found that enzyme networks exhibited a faster normalized response time when the *E_a_* of the bottleneck K_M_ was increased to >30 kcal/mol and the bottleneck enzymatic rate was greater than those of import and growth. Moreover, a TSEN composed of 5 bottlenecked intermediate reactions resulted in a more pronounced spike and deceleration (Fig. 7D), similar to our experimental measurements (Fig. 7C,D). The large spike and deceleration were due to rapid consumption of the network’s final metabolite (*c*_6_), which could only be replenished by upstream bottlenecked metabolites (Fig. 7E). The overall faster response time was captured by the model and was due to the decrease in steady-state concentration of *c*_6_at higher temperature caused by the large *E*_a_ of *K_M_* (Fig. 7E), leading *c*_6_ (which is limiting for growth) to stabilize more quickly.

With the same set of parameters, this TSEN model also predicted an under-shooting to below the new steady-state growth rate upon a temperature decrease from 32 °C to 22 °C (Fig. 7F), a nonintuitive behavior we confirmed experimentally (Fig. 7F). These findings suggest that the qualitatively distinct characteristics of the *S. pombe* temperature shift responses can be captured in a TSEN model by increasing the magnitude of *K_M_* parameters and the number of bottleneck reactions, and more generally that the growth network can be tuned to alter the balance between certain tradeoffs, to accelerate responses at the cost of temporary growth stalling.

## Discussion

Here, we used high-precision temperature control and single-cell analyses to show that bacteria exhibit a characteristic growth-rate response to temperature shifts within the Arrhenius range (Fig. 1C, 7A). The maintenance of the proteome across temperatures (Fig. 2) and the conservation of response dynamics during upshifts across mutants (Fig. 2G, 5E, S10I) and antibiotic treatments (Fig. 3H, S11B,D) collectively argue against the existence of a single regulator of growth rate across Arrhenius temperatures. Instead, the ability of our TSEN model to recapitulate nearly all observed behaviors (Fig. 5,6, S12) indicates that temperature sensitivity of growth is a collective property of the metabolic network, which encodes a memory of temperature states (Fig. 1I, S6, 6C). At steady state, the TSEN model predicts that growth depends only on substrate import kinetics and the total intracellular metabolite pool (Eq. 3), recovering the empirical Monod Equation (Eq. 4) and providing a simple framework that connects concentration-dependent growth directly to measured transporter kinetics^57^. Thus, our work establishes a mechanism for how Arrhenius-like growth arises across organisms^10^ (Fig. 1A, S1, S2, S12G).

A key feature is the temperature dependence of *K_M_*, which provides a straightforward mechanism for the asymmetry between upshift and downshift responses (Fig. 7F): if the substrate concentration of a reaction is near the *K_M_* at 27 °C (Fig. 7F, blue circle), an upshift to 37 °C will be accompanied by a small instantaneous growth rate increase since *K_M_* is higher at 37 °C versus 27 °C; vice versa, a downshift from the *K_M_* at 37 °C (Fig. 7F, red circle) will result in a larger instantaneous change in growth rate (Fig. 7F). Each of these instantaneous changes is then followed by an increase (upshift) or decrease (downshift) in the substrate concentration, and thus the growth rate (Fig. 7F), as predicted by the TSEN model (Fig. 5E, SI). The temperature upshift response is predicted to be largely insensitive to the *K_M_* of import at substrate saturation (Fig. 6F), suggesting that temperature-shift responses are independent of the details of import kinetics. A temperature-sensitive *K_M_* predicts that the *E_a_* of growth rate decreases with substrate concentration^12^ (Fig. 6H), consistent with our measurements of growth on amino acids (Fig. S13G). Such a decrease in temperature sensitivity at low substrate concentration is predicted by the TSEN to decrease the response time and increase the relative spike height (Fig. 6F,G,I,J), consistent with our experimental observations that decreases in amino acid concentrations resulted in faster responses to temperature upshifts and larger relative spikes (Fig. 6K,L). Thus, microorganisms confronting nutrient-poor environments may exhibit less temperature sensitivity of growth, somewhat paradoxically owing to the temperature sensitivity of *K_M_* values.

To produce the observed initial spike in growth rate, a bottleneck reaction with a large and highly temperature-sensitive *K_M_* was required (Fig. 5C). Previous measurements of intracellular metabolite dynamics in *E. coli* grown on glucose showed that reactions in central carbon metabolism are largely unsaturated and many reactions can exhibit large *K_M_* values (>1 mM)^59^ (Fig. 7G), indicating that the existence of such bottlenecks is highly likely. There are many potential bottleneck enzymes (Fig. 7G); those with the largest substrate concentrations are involved in aspartate consumption for amino acid (*aspC*) and pyrimidine (*pyrB*) biosynthesis, while those with the largest *K_M_* are involved in threonine (*ilvA*) and serine *(sdaA*, *ydfG)* degradation. As several of these reactions are essential, elucidating which reactions are true bottlenecks will likely require a combination of CRISPRi manipulation of gene expression, direct measurements of intracellular metabolite concentrations^59^, and biochemical characterization across temperatures.

Additionally, the TSEN model was able to explain qualitatively distinct temperature shift responses in the fission yeast *S. pombe*, which responded to an upshift with overshooting and deceleration (Fig. 7C), but which ultimately reached its steady-state growth rate faster than bacteria (Fig. 7D). The TSEN model predicted that large, coupled bottleneck reactions account for this behavior (Fig. 7D,E), and such a model predicted the undershoot response to a temperature downshift (Fig. 7F). In this scenario, faster temperature-shift responses come at the cost of increasing the activation energy of growth (Fig. 7C,F), further suggesting tradeoffs between fast steady-state growth across temperatures and the ability to respond quickly to shifts between temperatures.

Taken together, these findings suggest that metabolite flux rather than proteome rearrangement underlies growth rate adaptation across Arrhenius temperatures. As a result, cells adapt to temperature fluctuations without additional protein synthesis, resulting in transient memory of previously experienced temperatures (Fig. 2I, S7, 6C–E). Proteins comprise ∼55% of the cell’s biomass^62^ but constitute a relatively small total concentration ∼7 mM^59^ compared with metabolites, which make up ∼300 mM despite being only ∼3% of biomass^59,62^. Thus, increasing protein concentrations is metabolically expensive, and simulations of our TSEN model explicitly including protein synthesis confirmed that enzyme concentrations remain virtually constant throughout temperature shifts (Fig. S15). The conservation of growth rate responses across organisms (Fig. 7A–D), and the ability of the TSEN model to capture diverse responses to temperature shifts (Fig. 5–7) suggests an evolutionary pressure to adopt this strategy. The ability of *E. coli* cells to withstand fluctuations across a large range of temperatures, even short pulses at heat- and cold-shock temperatures (Fig. S7B,C,E), indicates a robustness that may be particularly important in the context of host colonization and infection-induced fevers, and our findings indicate that investigations of temperature adaptation can provide key insight into the metabolic factors limiting growth.

## Acknowledgements

We thank members of the Huang lab for helpful discussions. The *E. coli* BW25113 transposon mutant library was provided by the Deutschbauer lab, with help from Valentine Trotter. This work was funded by a Stanford Interdisciplinary Graduate Fellowship (to B.D.K.), NIH RM1 GM135102 (to K.C.H.) and NSF Awards EF-2125383 and IOS-2032985 (to K.C.H.). K.C.H. is a Chan Zuckerberg Biohub Investigator.

## Methods

### Culturing conditions and bacterial strains

*E. coli* wild-type and isolate strains were grown directly from 25% glycerol stocks stored at −80 °C in target media without selection. In typical temperature upshift experiments, cells were grown initially at 37 °C from frozen stocks overnight until saturation, then diluted 1:200 into fresh media at 37 °C until log phase (1.5–2 h in LB, ∼4 h in MOPS + glucose). Cells were then diluted 1:10 into fresh media and grown for at least 3 doublings at the lower, target temperature before performing a temperature upshift.

Most mutants (e.g., Keio collection knockouts) were grown from frozen stocks in target media with antibiotic selection. The ppGpp null strain (*spoT::cat, relA::kan*) was grown on LB plates with 10 µg/mL chloramphenicol and 25 µg/mL kanamycin overnight at 37 °C, and individual colonies were selected for further liquid culturing under selection to avoid suppressor mutations.

To evaluate growth kinetics on various media, *E. coli* MG1655 was first grown shaking overnight at 37 °C in MOPS minimal medium (Teknova, Hollister, CA) adjusted to pH 7.2 supplemented with 0.2% (w/v) D-glucose. To measure growth at lower temperatures (25 °C, 30 °C), cells were then diluted 1:200 in fresh MOPS+glucose and grown shaking until saturation for an initial passage at the target temperature. One milliliter of cells were washed in MOPS buffer (pH 7.2) at room temperature and 1 μL was added to 200 μL of target medium in 96-well plates for generating growth curves at the target temperature in a plate reader.

All strains used in this study are listed in Table S1.

### Liquid growth curves and analysis

Growth curves were measured using a protocol developed for accurately determining growth rates at low optical density^63^. Briefly, 200 μL of medium (without bacteria) were placed into each well of a transparent 96-well plate (Greiner Bio-One, Monroe, NC) and sealed with a transparent film (Excel Scientific, Victorville, CA) with holes for gas exchange cut above each well using a laser cutter (Epilog, Golden, CO). Optical density (OD) was measured with a BioTek Epoch 2 microplate spectrophotometer (Agilent, Santa Clara, CA) for at least 15 min to obtain blank values for each well at the target temperature. The seal was then removed, bacterial samples were added into each well, and the plate was sealed with a fresh laser-cut transparent film. Linear and orbital shaking were conducted between OD readings, which were taken every ∼7 min. The OD was corrected for non-linearity (linear range = 0–0.6) via a serial dilution of concentrated cells and performing a polynomial fit to obtain general fit parameters ^63^. For each well, the well-specific blank OD value determined prior to addition of cells was subtracted from the OD at each time point, which was then used to computed growth rate as a moving linear regression of the logarithm of the blanked OD.

### *E. coli* natural isolates

*E. coli* strains from non-human hosts were previously isolated^30,31,64^ from fecal samples collected from a variety of sources, including park and pet store animals and domestic pets, which were grown on Colilert-18 medium (IDEXX, Portland, ME) for selection of presumptive *E. coli* colonies^31^. Isolate identities were confirmed by beta-glucuronidase activity and subsequent sequencing of the corresponding gene, *uidA*^31^. The strains were grown overnight in a rich medium (LB) at 37 °C, diluted 1:200 in fresh LB, and grown for 24 h across temperatures from 27 °C–47 °C to measure activation energies.

### Temperature-controlled single-cell imaging

The temperature control platform, the <u>Si</u>ngle-<u>C</u>ell <u>Te</u>mperature <u>C</u>ontroller (SiCTeC), was designed and described previously^32^. Briefly, a ring-shaped Peltier module (TE Technology, Traverse City, MI) was adhered to a glass slide, and the sample temperature was monitored on the coverslip and controlled using a micro-Arduino with a proportional-integral-derivative (PID) algorithm. Sample temperature was monitored and visualized in real time using the open-source software Processing^65^. Agarose hydrogels were prepared by boiling 3% ultrapure agarose (Sigma Aldrich, St. Louis, MO) in the target medium, and 200 μL of the mixture were pipetted onto a 9-mm diameter silicone gasket (Grace Bio-Labs, Bend, OR) onto the temperature-controlled glass slide. An additional slide was placed on the gasket to compress the hydrogel, which then cooled and solidified at the initial temperature of the experiment. After removing the additional compressing slide, 1 μL of cells was pipetted onto the solidified hydrogel and dried briefly (<1 min) at the initial temperature of the experiment.

Imaging was performed on a Ti-Eclipse microscope in phase-contrast mode using a 40X Ph2 air objective (NA: 0.95) (Nikon Instruments, Tokyo, Japan) with a 1.5X tube lens. The air objective was used to avoid heatsink issues with oil-immersion objectives. Images were captured every 30 s on a Zyla 4.2 sCMOS (Andor Technology, Belfast, UK), Neo 5.5 sCMOS (Andor Technology, Belfast, UK) or PCO Panda 4.2 (Excelitas, Pleasanton, CA) scientific camera. The microscope system was integrated using μManager 1.41^66^.

### Image analysis

To extract cell morphology information throughout the experiment, subpixel-resolution cellular contours were obtained through a combination of deep learning-based and traditional image segmentation software^32,67^. Briefly, we first aligned the images using the Template-Matching plugin in FIJI^68,69^, then each image was processed with a fully convolutional neural network model, *DeepCell*^67^. Separate neural networks were trained for *E. coli* rich medium, *E. coli* minimal medium, and *S. pombe* rich medium, with >200 cells manually annotated in each condition^32,67,70^. Outputs from DeepCell classification were used to extract cellular contours using *Morphometrics* v. 1.1 in MATLAB (MathWorks, Natick, MA)^71^. Custom MATLAB scripts were used to track individual cells and measure cellular geometry^70^.

### Rapid temperature downshifts

To increase the rate of temperature change during downshifts from previous experiments performed using ambient temperature as the cooling sink, which required ∼5 min from 37°C to 27 °C^32^, we used dry ice (solid CO_2_) to rapidly cool samples. During a downshift, the SiCTeC device was powered off and a 250-mL beaker with dry ice was used to pour sublimating dry ice directly onto the sample on the glass slide. For downshifts to temperatures above ambient, when the temperature was ∼1 °C above the target downshift temperature, the SiCTeC device was powered back on and the PID algorithm restarted control of the sample temperature. The sample temperature was closely monitored and adjusted accordingly for undershooting. This method enabled stabiliziation at the lower temperature within <1 min. For downshifts to temperatures below ambient, dry ice was continuously poured onto the sample at intervals necessary to maintain the target temperature. Temperature upshifts back to 37 °C were performed by turning back on the SiCTeC device.

### Anaerobic growth

To perform temperature shifts in anerobic conditions, *E. coli* MG1655 was inoculated from a frozen stock into LB and grown overnight in an anaerobic chamber (Coy Laboratory Products, Grass Lake, MI), then diluted 1:200 and grown until saturation in pre-reduced LB (i.e., kept in chamber for >48 h before culturing). Cells were diluted 1:200 in pre-reduced LB and grown at 37 °C until log phase (∼2 h), then diluted 1:10 and grown at 27 °C for 3 h. From log phase at 27 °C, 1 μL was pipetted on a SiCTeC glass slide for temperature shifts (all components pre-reduced) and imaged inside the chamber using a Ti-Eclipse inverted microscope (Nikon Instruments, Tokyo, Japan) with a Neo 5.5 sCMOS (Andor Technology, Belfast, UK). The SiCTeC platform was controlled by a laptop inside the chamber.

### Quantification of response time

To measure the response time of an upshift trajectory, the steady-state growth rate (*g^ss^*) at the final temperature was determined by measuring the average of the growth rate across the timepoints after which growth rate had stabilized (determined by visual inspection). Its error (*δg^ss^*) is defined as the standard error of the mean. This steady-state growth rate was subtracted from the entire trajectory (*g*(*t*) – *g^ss^*), and the response time was determined as the time (*t*) at which *g*(*t*) was indistinguishable from *g^ss^* within error (*δg^ss^*), which was generally ∼5% of *g^ss^*.

### Antibiotic treatment during single-cell temperature shifts

Five-hundred microliter aliquots of growth medium+3% ultrapure agarose were melted at 95 °C, then mixed with 0.5 μL of 1000X target concentration antibiotic stock solution and 150 μL was immediately used to make a hydrogel for the sample as described above. The short time exposed to higher temperatures (∼30 s) likely only moderately impacted the minimum inhibitory concentration, as antibiotic efficacies against *E. coli* are unaffected by long-term (30 min) treatment at 56 °C, with many drugs unaffected by autoclaving (121 °C)^72^.

### Liquid culture samples for proteome extraction

*E. coli* MG1655 was grown at 37 °C overnight in MOPS buffer (Teknova, Hollister, CA) adjusted to pH 7.2 supplemented with 0.2% (w/v) of carbon source (glucose or glycerol) (Sigma-Aldrich, St. Louis, MO) or LB (Thermo Fisher Scientific, Waltham, MA), then diluted in duplicate and grown for at least 4 doublings at the target temperature (25 °C, 30 °C, or 37 °C). Growth at additional temperatures (16 °C, 43 °C) were assayed for LB. Cultures were grown to log phase (OD_600_∼0.2) in 50-mL Falcon tubes and 15 mL were harvested and washed with 1 mL PBS via centrifugation at 4 °C. Supernatant was removed and the pellet was snap-frozen with liquid nitrogen.

### Proteome extraction

Extraction was performed as previously described^73^. Briefly, samples were thawed and lysed using a bead-beating procedure, then supernatant was reduced and alkylated with DTT and iodoacetamide, respectively. Peptides were then washed, digested, and eluted using S-trap tubes (Protifi) and desalted with C18 solid-phase extraction (Sep-Pak Waters). Finally, peptides were dried by vacuum centrifugation and quantified for normalization (Nanodrop ND-1000).

### Proteomic analysis via LC-MS/MS and database searching

Peptide quantification was performed as previously described^73^. Briefly, dried peptides were diluted in 0.2% formic acid to a final concentration of 0.5 μg/mL, then separated by reversed-phase chromatography (Dionex Ultimate 3000 HPLC, Thermo Fisher Scientific, Waltham, MA) and directed to a LTQ Orbitrap Elite mass spectrometer (Thermo Fisher Scientific, Waltham, MA). The top 20 ions were selected for fragmentation using collision-induced dissociation (DIC), and fragments were analyzed in the ion trap with rapid scan rate.

Mass spectra were searched against the UniProt canonical *E. coli* FASTA database using the SEQUEST algorithm of Proteome Discoverer 2.2.0.388. Peptides were filtered using Percolator with the protein false discovery rate set to 1%. Protein abundance was based on precursor ion peak areas.

### Proteome data analysis and annotation using Clusters of Orthologous Groups (COG)

Protein functional annotation was performed using the Clusters of Orthologous Groups of proteins (COG) database^42^. UniProt accession codes were mapped to the COG database downloaded from https://ftp.ncbi.nih.gov/pub/COG/COG2014/data/. After removing non-bacterial groups, COGs were reduced to 9 functional groups: ribosomal protein (RP), non-RP translational, transcription, DNA replication, cell division, cell envelope structure, post-translational modification, energy and metabolism, and other. The corresponding gene of each protein was annotated using the gene association table from EcoCyc (https://ecoliwiki.org/gaf/gene_association.ecocyc.gz)^74^. Further analysis was performed using custom MATLAB scripts, wherein the relative abundance of each protein was calculated for each sample and its replicates and then compared across conditions.

### *E. coli* transposon library temperature shifts

One milliliter of a pooled library of *E. coli* BW25113 transposon insertion mutants with random DNA barcodes^44^ was thawed and diluted into 50 mL of LB+25 μg/mL kanamycin (Sigma Aldrich, St. Louis, MO) in a 250-mL Erlenmeyer flask (Pyrex, Glendale, AZ) and grown shaking overnight at 37 °C. 0.5 mL of the stationary-phase library cultures were diluted into 50 mL of LB in technical replicates at 25 °C and 37 °C.

The 25 °C culture was grown for 3 h to early log phase (OD_600_∼0.15) and 6 mL were diluted into in each of two flasks containing 50 mL fresh LB pre-warmed at 25 °C, then grown for another 3 h in an air-heated shaker (New Brunswick Scientific, Edison, NJ). The temperature upshift from 25 °C to 37 °C was performed by placing one of the 25 °C flasks into a heated water shaker at 37 °C. The control sample was left at 25 °C. Samples were collected in 1.5-mL aliquots and flash-frozen using liquid nitrogen every 10 min at timepoints of −10, 0, 10, 20, 30, 40, 60, 90 min relative to the shift (control samples without temperature shifts were also taken at the same time points).

The 37 °C culture was grown for 2 h until mid- to late-log phase (OD_600_∼0.3) and 1.5 mL were diluted into each of two flasks containing 50 mL of fresh LB pre-warmed at 37 °C, then grown for ∼1 h in a heated water shaker. For the temperature downshift from 37 °C to 25 °C, one of the flasks was first placed into a room-temperature water bath for 6 min for faster cooling, then transferred to an air-heated shaker at 25 °C. The other flask was maintained at 37 °C as a control.

Each set of downshift/upshift experiments was performed twice, and all samples were flash frozen and immediately stored at −80 °C. Optical density was monitored at each time point for all experiments with a Genesys 20 spectrophotometer (Thermo Fisher Scientific, Waltham, MA). The time scale of the temperature upshift was estimated to be ∼3.5 min using a virtual experiment with 50 mL of water in a flask, whose temperature was directly monitored during a shift from 25 °C to 37 °C using the SiCTeC device thermistor reading. A similar test was performed for the 37 °C to 25 °C downshift by placing a 37 °C flask with 50 mL of water into a stationary water bath at room temperature, giving an estimate of ∼6 min to complete the temperature shift.

### *E. coli* transposon mutant library sequencing and analysis

Barcode sequencing (BarSeq) was performed as previously described^44^. Briefly, genomic DNA was extracted using the DNeasy 96 Blood and Tissue kit (Qiagen) and quantified using the Quant-iT dsDNA BR Assay kit (Qiagen). BarSeq PCR was performed using 200 ng of genomic DNA template in 150-μL reaction volumes with Q5 polymerase (with enhancer) (New England Biolabs, Ipswich, MA), 20 μM universal forward P5 primer, and 20 μM reverse P7 primers with 6-bp TruSeq indices that are automatically demultiplexed by Illumina software. PCR products were checked for completion using gel electrophoresis, and 10 μL of each reaction product were pooled and purified using the Zymo DNA Clean and Concentrator kit (Zymo Research, Irvine, CA). Sequencing was performed at the Chan Zuckerberg BioHub facility on an Illumina NextSeq 550 platform in High Output mode.

BarSeq analysis was performed as previously detailed^44^, with relevant scripts at https://genomics.lbl.gov/supplemental/rbarseq/. Briefly, barcode reads were mapped to their corresponding genomic loci using sequencing of the transposon insertions in the *E. coli* library, and genes were filtered for those with ≥10 barcodes in the central (10–90%) portion of the gene. Temperature-shift samples were analyzed along with their corresponding unshifted control samples, with the time-zero sample corresponding to when the library was shifted (*t*=0 sample). The fitness of each mutant was measured as the log_8_ fold-change in its relative abundance, and further analysis was performed using custom MATLAB scripts.

### FRAP measurements

*E. coli* cultures were grown overnight from frozen stocks at 37 °C in LB, with 25 µg/mL kanamycin selection for Δ*fabR* and Δ*fadR* mutants, and then diluted 1:200 into fresh LB and grown until log phase at 37 °C (∼2 h). Cultures were then diluted 1:10 into fresh LB, and 2 µL of 0.5 mg/mL MitoTracker Green (Thermo Fisher Scientific, Waltham, MA) were added. The culture was grown at the target temperature for 30 min to enable sufficient membrane labeling. Five hundred microliters of stained cells were washed via centrifugation (30 s at 7000 rcf) in fresh LB at the target temperature (37 °C or 27 °C). One microliter of washed culture was placed on an LB 3% agarose hydrogel and prepared for imaging using standard glass slide technique. Imaging was performed with a Zeiss LSM 880 confocal microscope with an environmental chamber for monitoring and controlling temperature (37 °C) and integrated with the ZEN software suite (Zeiss, Oberkochen, Germany). Temperature upshift experiments were performed by placing the sample into the heated environmental chamber for 5 min before imaging. Fluorescence recovery after photobleaching (FRAP) of individual cells was performed with excitation at 488 nm using the ZEN software by choosing a photobleaching region near cell tips covering 1/4 to 1/3 of the cell. The entire cell was imaged at minimum frame intervals (150–300 ms) to image fluorescence recovery.

### FRAP analysis and viscosity estimate

Individual regions of interest were manually delineated for the photobleached cell tips and the entire cell in FIJI^69^, and the total fluorescence of each area was quantified at each time point. These data were imported for further analysis in MATLAB. The fluorescence recovery in each photobleached region was corrected by the rate of photobleaching in the entire cell, and the recovery curve was normalized and fit to the exponential function 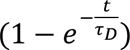. The time constant (*τ*_D_) was used to estimate a diffusion coefficient (*D*) via solution to Gaussian photobleaching in a plane, 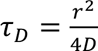 (i.e., modeling the cell tip as a disk of radius *r* on the membrane)^75^. The viscosity (η) was calculated using the Stokes-Einstein equation 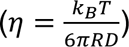, where *R* is the radius of the fluorophore (estimated as 2 nm).

### Schizosaccharomyces pombe culturing

*S. pombe* WT972 h-was grown from a frozen stock on YE5S plates (5 g/L yeast extract, 30 g/L glucose, 225 mg/L each of adenine, histidine, leucine, uracil, and lysine hydrochloride, 2% Difco Bacto Agar). A single colony was grown at 22 °C or 32 °C in liquid YE5S overnight until saturation, then diluted 1:100 and grown until log phase (OD_600_∼0.3–0.5). One microliter of log-phase cells was placed on a 3% agarose YE5S pad at the initial temperature of the experiment for imaging.

### Estimate of LB molarity

LB is largely comprised of free amino acids, and measurements of *E. coli* auxotrophy and direct HPLC quantification^74^ have provided an estimate of ∼122 g/mol for its free amino acid content. The molar mass of tryptone is reported by the manufacturer to be 71.08 g/mol (Thermo Fisher Scientific, Waltham, MA). As LB is composed of 10 g tryptone and 5 g yeast extract dissolved in 1 L H_2_O, we estimate that the concentration of metabolizable free amino acids in LB is ∼182 mM. Note that the accuracy of this estimate is not critical for any of our conclusions; the value simply enables plotting LB concentrations on the same plot as other substrates such as casamino acids.

### Measurements of K_M_

*E. coli* MG1655 was grown overnight until saturation in MOPS buffer+0.2% (w/v) glucose at the target temperature (25 °C, 30 °C, 37 °C), then washed twice in MOPS buffer before being diluted 1:200 in the target medium at the target temperature. Liquid culture growth curves were obtained as described above. Optical density (OD) was measured using a microplate reader and maximal growth rate was quantified as the peak of the derivative of ln(background-subtracted OD)^63^. A weighted fit was performed on the growth rate versus concentration curve using the Monod equation (Eq. 4) to extract an estimate of *K_M_* with a standard error. We note that it was challenging to obtain growth rates at low concentration for simple sugars (e.g., glucose, fructose), due to both the low *K_M_* and generally low growth rate (<0.2 h^−1^) (Fig. S13A–C). In succinate, the relationship between ln(growth rate) and 1/T appeared bilinear, thus a linear fit produced an estimate of *E_α_* with very large standard error (19±14 kcal/mol) (Fig. S13G).

