## Supplemental Information for "Metabolomic rearrangement controls the intrinsic microbial response to temperature changes"

**Supplementary Information for "Metabolomic rearrangement controls the intrinsic microbial response to temperature changes"**

**Authors:** Benjamin D. Knapp[1], Lisa Willis[2], Carlos Gonzalez[3], Harsh Vashistha[4], Joanna Jammal Touma[4], Mikhail Tikhonov[5], Jeffrey Ram[6], Hanna Salman[4], Josh E. Elias[7], Kerwyn Casey Huang[1,2,7,8,†]

**Affiliations:**

[1]Biophysics Program, Stanford University, Stanford, CA 94305, USA

[2]Department of Bioengineering, Stanford University, Stanford, CA 94305, USA

[3]Department of Chemical and Systems Biology, Stanford University School of Medicine, Stanford, CA 94305, USA

[4]Department of Physics and Astronomy, University of Pittsburgh, Pittsburgh, PA 15260, USA

[5]Department of Physics, Washington University in St. Louis, St. Louis, MO 63130, USA

[6]Department of Physiology, Wayne State University, Detroit, MI 48201, USA

[7]Chan Zuckerberg Biohub, San Francisco, CA 94158, USA

[8]Department of Microbiology and Immunology, Stanford University School of Medicine, Stanford, CA 94305, USA

### SUPPLEMENTARY TABLES

**Table S1: Strains in this study.**

| Organism | Genotype | Source and Reference |
|---|---|---|
| *Escherichia coli* | MG1655 (wild type) | Laboratory collection |
| *Escherichia coli* | BW25113 (wild type) | Laboratory collection |
| *Escherichia coli* | CS109 (wild type) | Kevin Young |
| *Escherichia coli* | BL21 (wild type) | Laboratory collection |
| *Escherichia coli* | NILS1 (human isolate) | Ref. [1] |
| *Escherichia coli* | Ram17 (pig isolate) | Jeffrey Ram |
| *Escherichia coli* | Ram146 (raccoon isolate) | Jeffrey Ram |
| *Escherichia coli* | Ram376 (snake isolate) | Jeffrey Ram |
| *Escherichia coli* | Ram551 (turtle isolate) | Jeffrey Ram |
| *Escherichia coli* | Ram568 (turtle isolate) | Jeffrey Ram |
| *Escherichia coli* | Ram569 (tutle isolate) | Jeffrey Ram |
| *Escherichia coli* | Ram581 (goose isolate) | Jeffrey Ram |
| *Escherichia coli* | Ram589 (seagull isolate) | Jeffrey Ram |
| *Escherichia coli* | Ram592 (pigeon isolate) | Jeffrey Ram |
| *Escherichia coli* | Ram1584 (dog isolate) | Jeffrey Ram |
| *Escherichia coli* | Ram1612 (human isolate) | Jeffrey Ram |
| *Escherichia coli* | Ram1636 (horse isolate) | Jeffrey Ram |
| *Escherichia coli* | Ram1897 (seagull isolate) | Jeffrey Ram |
| *Escherichia coli* | *dnaK::kan* | Keio collection |
| *Escherichia coli* | *stpA::kan* | Keio collection |

| | | |
|---|---|---|
| *Escherichia coli* | *sodB::kan* | Keio collection |
| *Escherichia coli* | *nemA::kan* | Keio collection |
| *Escherichia coli* | *ompT::kan* | Keio collection |
| *Escherichia coli* | *fabR::kan* | Keio collection |
| *Escherichia coli* | *fadR::kan* | Keio collection |
| *Escherichia coli* | *uup::kan* | Keio collection |
| *Escherichia coli* | *ettA::kan* | Keio collection |
| *Escherichia coli* | *ybiT::kan* | Keio collection |
| *Escherichia coli* | *yheS::kan* | Keio collection |
| *Escherichia coli* | *bipA::kan* | Keio collection |
| *Escherichia coli* | *lepA::kan* | Keio collection |
| *Escherichia coli* | *clpP::kan* | Keio collection |
| *Escherichia coli* | *lon::kan* | Keio collection |
| *Escherichia coli* | *tusA::kan* | Keio collection |
| *Escherichia coli* | *tusB::kan* | Keio collection |
| *Escherichia coli* | *relA::kan spoT::cat* (MG1655 parent) | Ref. [2] |
| *Bacillus subtilis* | 168 (wild-type) | Laboratory collection |
| *Escherichia fergusonii* | B253 wild-type | Laboratory collection |
| *Schizosaccharomyces pombe* | 972 h- (wild-type) | Laboratory collection |

24

### SUPPLEMENTARY FIGURES

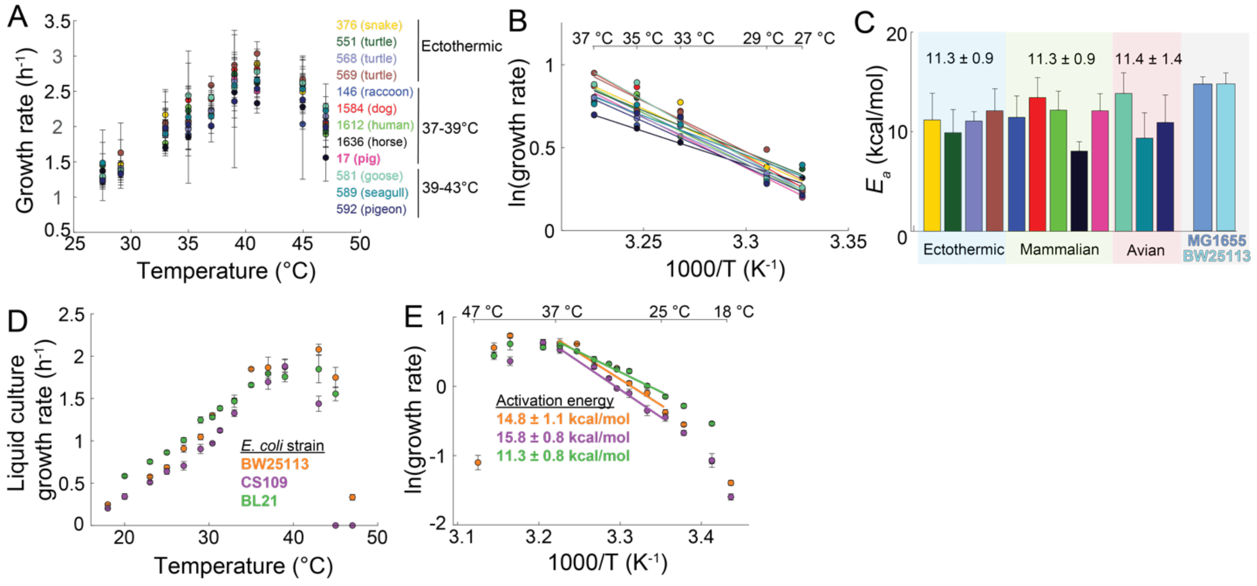


**Figure S1: *E. coli* exhibits robust Arrhenius behavior with a highly conserved activation energy.**

A) Liquid-culture maximal growth rates across temperatures of diverse *E. coli* strains from various animal hosts (Methods). Each strain is labeled with its laboratory accession number (Table S1), host source, and estimated host body temperature.

B) Arrhenius plots of growth rates from (A). The natural logarithm of maximal growth rate is plotted against the inverse absolute temperature for temperatures between 27 °C and 37 °C, along with weighted linear fits for each strain.

C) Activation energies measured as the slope of the linear fit to the data in (B) for each *E. coli* strain, with errors reported as the standard error of the mean (SEM) from the weighted fit. Each strain is grouped according to host body temperature (blue: Ectothermic, green: Mammalian, red: Avian, gray: Laboratory (MG1655, BW25113)).

D) Steady-state maximum growth rates in rich medium (LB) of *E. coli* BW25113, CS109, and BL21 between 18 °C and 47 °C (Methods). Each maximal growth rate is reported as the mean±1 standard deviation of >6 replicates.

E) Arrhenius plot of growth rates from (D). The natural logarithm of maximal growth rate is plotted against the inverse absolute temperature. Growth rates measured at temperatures between 25 °C and 37 °C were used for measuring the activation energy (slope, $E_a$).

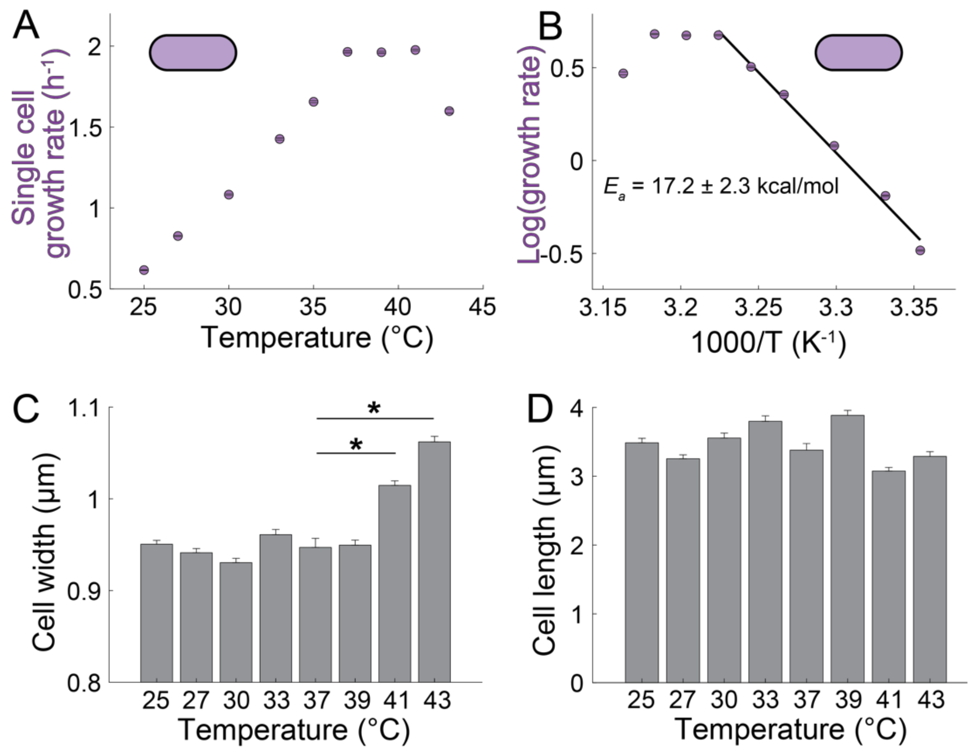


**Figure S2: *E. coli* exhibits single-cell Arrhenius behavior with highly conserved cell shape.**

A) Steady-state single-cell growth rates of *E. coli* MG1655 grown on rich medium (LB) between 25 °C and 43 °C (Methods). Each growth rate is reported as the mean±1 SEM (error bars not visible). Instantaneous growth rates were measured from *n*=97-414 cells in each experiment.

B) Arrhenius plot of single-cell growth rates from (A). The natural logarithm of steady-state growth rate is plotted against the inverse absolute temperature. Growth rates measured at temperatures between 25 °C and 37 °C were used for measuring the activation energy (slope, $E_a$).

C) Width of *E. coli* cells during steady-state growth on rich medium (LB) between 25 °C and 43 °C. Cell width distributions were significantly different between 37 °C and elevated temperatures (41, 43 °C). Significance was determined via a two-sample *t-test*. *: $p<0.05$. Values are reported as mean±1 SEM. $n$=97-414 cells in each experiment.

D) Length of *E. coli* cells during steady-state growth on rich medium (LB) between 25 °C and 43 °C. Cell length distributions were not significantly different between any two temperatures. Significance was determined via a two-sample *t-test*. Values are reported as mean±1 SEM. $n$=97-414 cells in each experiment.

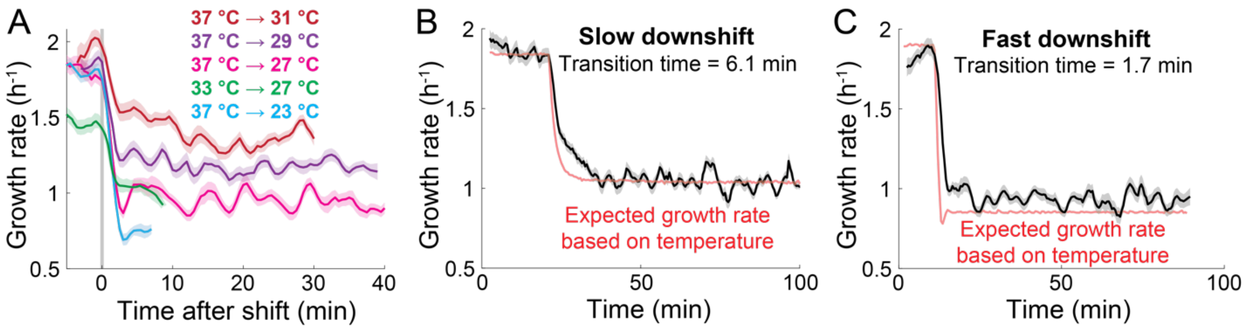


**Figure S3: Responses to temperature downshifts follow the downshift time scale.**

A) Single-cell growth rates of *E. coli* MG1655 on rich medium (LB) undergoing temperature downshifts between 37 °C and 23 °C. The growth rate is reported as the mean±1 SEM (shaded region) at each time point. *n*=192-412 cells in each experiment.

B) Single-cell growth rates (black) of *E. coli* MG1655 on rich medium (LB) undergoing a slow temperature downshift from 37 °C to 27 °C. The expected growth rate (red) was generated by fitting the temperature data to an Arrhenius function with parameters set by the initial and final growth rates of the trajectory. *n*=762 cells.

C) Single-cell growth rates (black) of *E. coli* MG1655 on rich medium (LB) undergoing a fast temperature downshift from 37 °C to 27 °C. The expected growth rate (red) was generated by fitting the temperature data to an Arrhenius function with parameters set by the initial and final growth rates of the trajectory. *n*=192 cells.

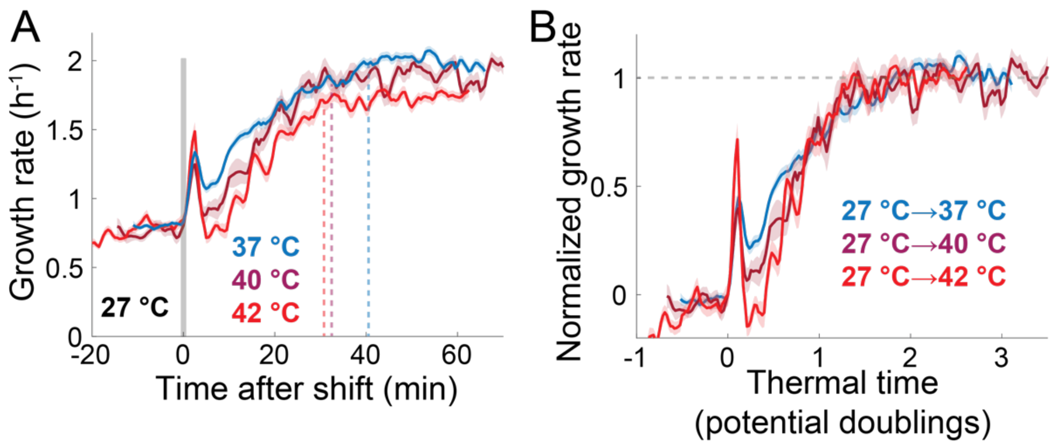


**Figure S4: Temperature upshifts to mild heat-shock temperatures are characterized by slower final growth rates but similar normalized response times.**

A) Single-cell growth rates of *E. coli* MG1655 on rich medium (LB) undergoing temperature upshifts from 27 °C to 37 °C (blue, *n*=792 cells), 40 °C (purple, *n*=278 cells), or 42 °C (red, *n*=474 cells). Data are the mean±1 SEM (shaded region) at each time point.

B) Normalized growth rate versus thermal time follows a common trajectory for each upshift in (A).

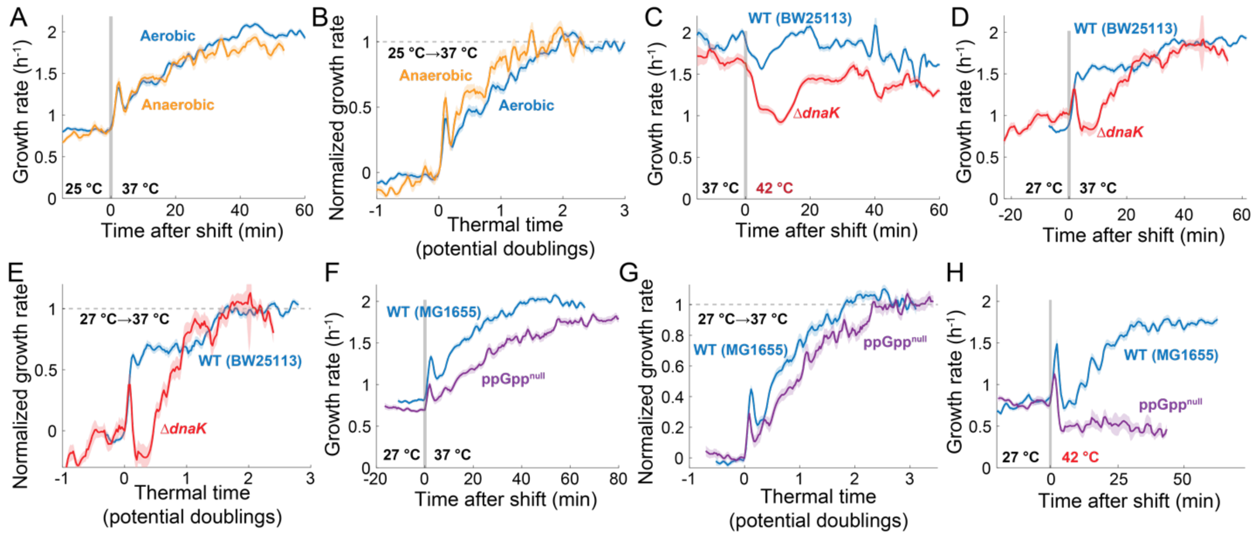


**Figure S5: Effect of chaperones, oxygen, and the stringent response on temperature upshift responses.**

A) Single-cell growth rates of *E. coli* MG1655 on rich medium (LB) undergoing a temperature upshift from 25 °C to 37 °C in aerobic (blue, *n*=773 cells) or anaerobic (orange, *n*=319 cells) conditions. Data are the mean±1 SEM (shaded region) at each time point.

B) Normalized growth rate versus thermal time for each trajectory in (A).

C) Single-cell growth rates of Δ*dnaK* (red, *n*=433 cells) and its parent BW25113 (blue, *n*=1011 cells) on rich medium (LB) undergoing a temperature upshift from 37 °C to 42 °C. Data are the mean±1 SEM (shaded region) at each time point.

D) Single-cell growth rates of Δ*dnaK* (red, *n*=318 cells) and its parent BW25113 (blue, *n*=734 cells) on rich medium (LB) undergoing a temperature upshift from 27 °C to 37 °C. Data are the mean±1 SEM (shaded region) at each time point.

E) Normalized growth rate versus thermal time for each trajectory in (D).

F) Single-cell growth rates of a ppGpp$^{null}$ strain (Δ*relA* Δ*spoT*) (purple, *n*=648 cells) and its parent MG1655 (blue, *n*=792 cells) on rich medium (LB) undergoing a temperature upshift from 27°C to 37 °C. Data are the mean±1 SEM (shaded region) at each time point.

G) Normalized growth rate versus thermal time for each trajectory in (F).

H) Single-cell growth rates of ppGpp$^{null}$ (purple, *n*=47 cells) and its parent MG1655 (blue, *n*=474 cells) on rich medium (LB) undergoing a temperature upshift from 27°C to 42 °C. Data are the mean±1 SEM (shaded region) at each time point.

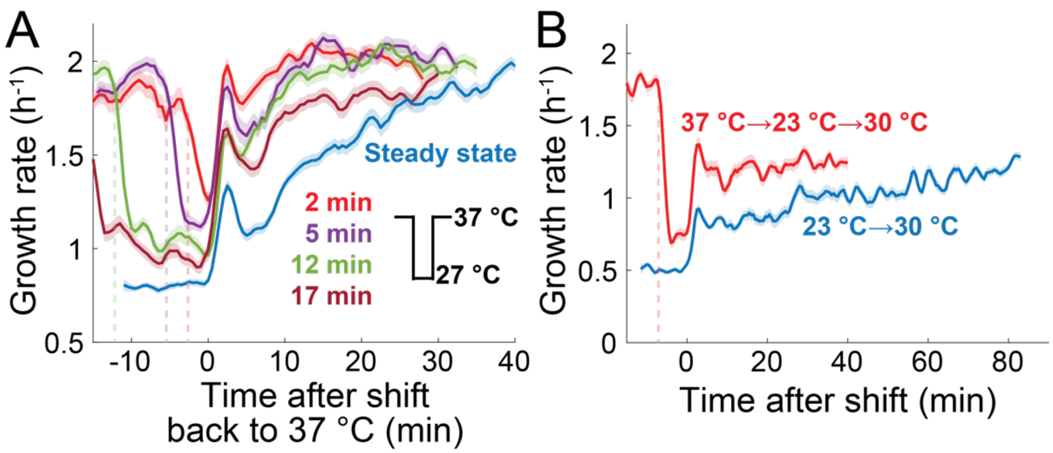


**Figure S6: Downshift pulses reveal temperature history.**

A) Single-cell growth rates of *E. coli* MG1655 on rich medium (LB) starting at 37 °C subjected to 27 °C pulses for 2 min (red, *n*=519 cells), 5 min (purple, *n*=397 cells), 12 min (green, *n*=451 cells), or 17 min (dark red, *n*=422 cells). Dotted lines represent the time at which cells were subjected to a 27 °C downshift. The shift from steady-state growth at 27 °C to 37 °C is also shown for comparison (blue, *n*=773 cells). Data are the mean±1 SEM (shaded region).

B) Single-cell growth rates of *E. coli* MG1655 on rich medium (LB) starting at 37 °C subjected to a 23 °C pulse for 5 min before an upshift to the intermediate temperature 30 °C (red, *n*=330 cells). The vertical dashed line indicates the start of cooling, which required ~2 min to reach 23 °C. The shift from steady-state growth at 23 °C to 37 °C is also shown for comparison (blue, *n*=396 cells).

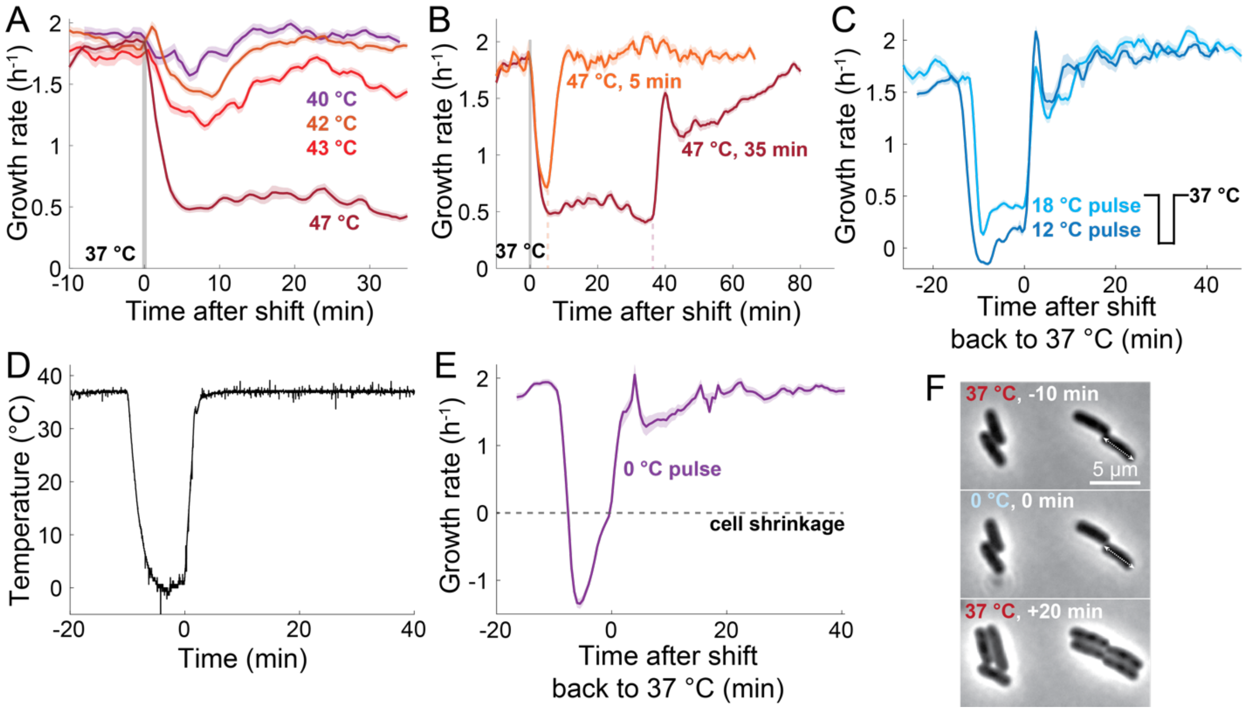


**Figure S7: *E. coli* growth rate responds rapidly to heat-shock and cold-shock pulses.**

A) Single-cell growth rates of *E. coli* MG1655 on rich medium (LB) undergoing temperature upshifts from 37 °C to 40 °C (purple, *n*=479 cells), 42 °C (orange, *n*=1249 cells), 43 °C (red, *n*=819 cells), or 47 °C (dark red, *n*=474 cells). Data are the mean±1 SEM (shaded region) at each time point.

B) Single-cell growth rates of *E. coli* MG1655 on rich medium (LB) starting at 37 °C and subjected to heat-shock pulses at 47 °C for 5 min (orange, *n*=381 cells) or 35 min (dark red, *n*=474 cells). Vertical dashed lines represent the times at which cells were shifted back to 37 °C. Data are the mean±1 SEM (shaded region) at each time point.

C) Single-cell growth rates of *E. coli* MG1655 on rich medium (LB) starting at 37 °C and subjected to ~10-min cold-shock pulses at 18 °C (light blue, *n*=361 cells) or

12 °C (dark blue, *n*=390 cells). Data are the mean±1 SEM (shaded region) at each time point.

D) Temperature readout of a 5-min pulse at 0 °C (Methods) starting from 37 °C. *t*=0 is when cells were shifted back to 37 °C.

E) Single-cell growth rates of *E. coli* MG1655 on rich medium (LB) starting at 37 °C and subjected to a 5-min cold-shock pulse at 0 °C (purple, *n*=283 cells) shown in (D). Horizontal dashed line represents cell shrinkage defined as growth rate <0 $h^{-1}$. Data are the mean±1 SEM (shaded region) at each time point.

F) Images of *E. coli* MG1655 on rich medium (LB) during a 10-min pulse at 0 °C starting from 37 °C. At 37 °C, cells exhibited normal morphologies and growth (top). At 0 °C, cells shrank, as exemplified by the cell whose length at *t*=-10 min represented by a double-arrowed line extends beyond the cell boundary at *t*=0 (middle). Growth resumed quickly after the sample was heated back to 37 °C, without loss in cell viability (bottom).

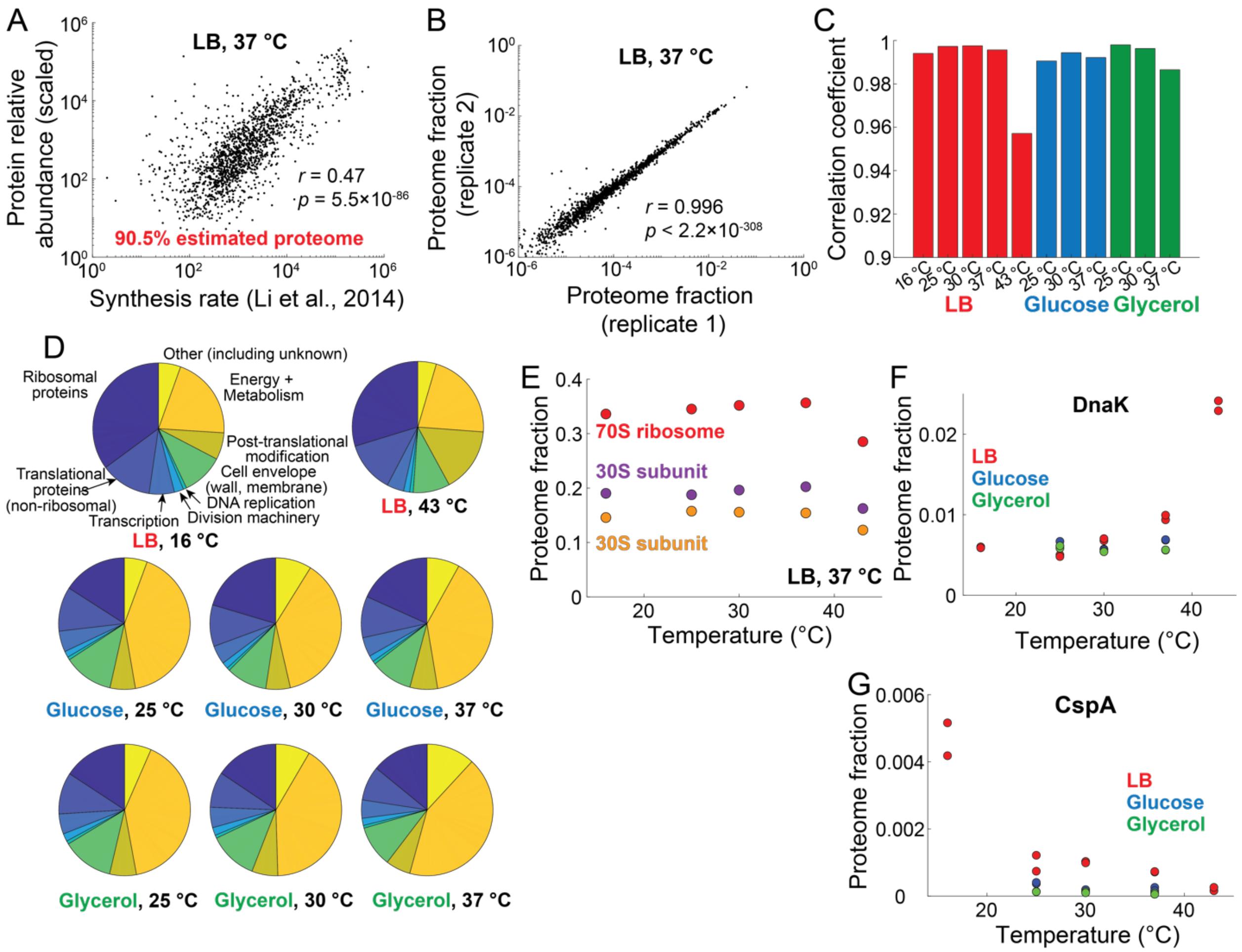


**Figure S8: Proteomics reveals a temperature-independent proteome in *E. coli*.**

A) Mean relative abundance of proteins from 37 °C rich medium (LB) samples measured by LC-MS/MS scaled by estimated total protein count from ribosomal profiling versus absolute abundance estimate from ribosome profiling in Ref. [3]. The datasets are in good agreement, and the proteomics dataset recovers 90.5% of the proteome.

B) Correlation of protein relative abundance measurements between biological replicates of 37 °C rich medium (LB) samples. Each dot represents a single protein (*n*=1918).

C) Correlation coefficients between biological replicates were high for all proteomics samples. Coefficients are ordered by growth medium (LB, glucose, glycerol) and then by temperature.

D) The *E. coli* proteome at 16 °C and 43 °C in rich medium (LB) (top row), and the proteome at 25 °C, 30 °C, and 37 °C in glucose (middle) and glycerol (bottom). Functional proteomic sectors are annotated according to a modification of the Clusters of Orthologous Groups (Methods).

E) Proteome fraction represented by ribosomal subunits across all temperatures in rich medium (LB). Fractions for the small subunit (30S, orange), large subunit (50S, purple), and full ribosome (70S, red) did not significantly vary between 16 °C and 37 °C. Values are the mean of biological replicates.

F) Proteome fraction represented by the protein DnaK versus temperature for each sample, colored by growth medium (LB, glucose, glycerol). Replicates are shown as individual filled circles. At 43 °C, DnaK increased substantially to >2% of the proteome.

G) Proteome fraction represented by the protein CspA versus temperature for each sample, colored by growth medium (LB, glucose, glycerol). Replicates are shown as individual filled circles. CspA increased from ~0.1% of the proteome at temperatures ≥25 °C to ~0.5% at 16 °C.

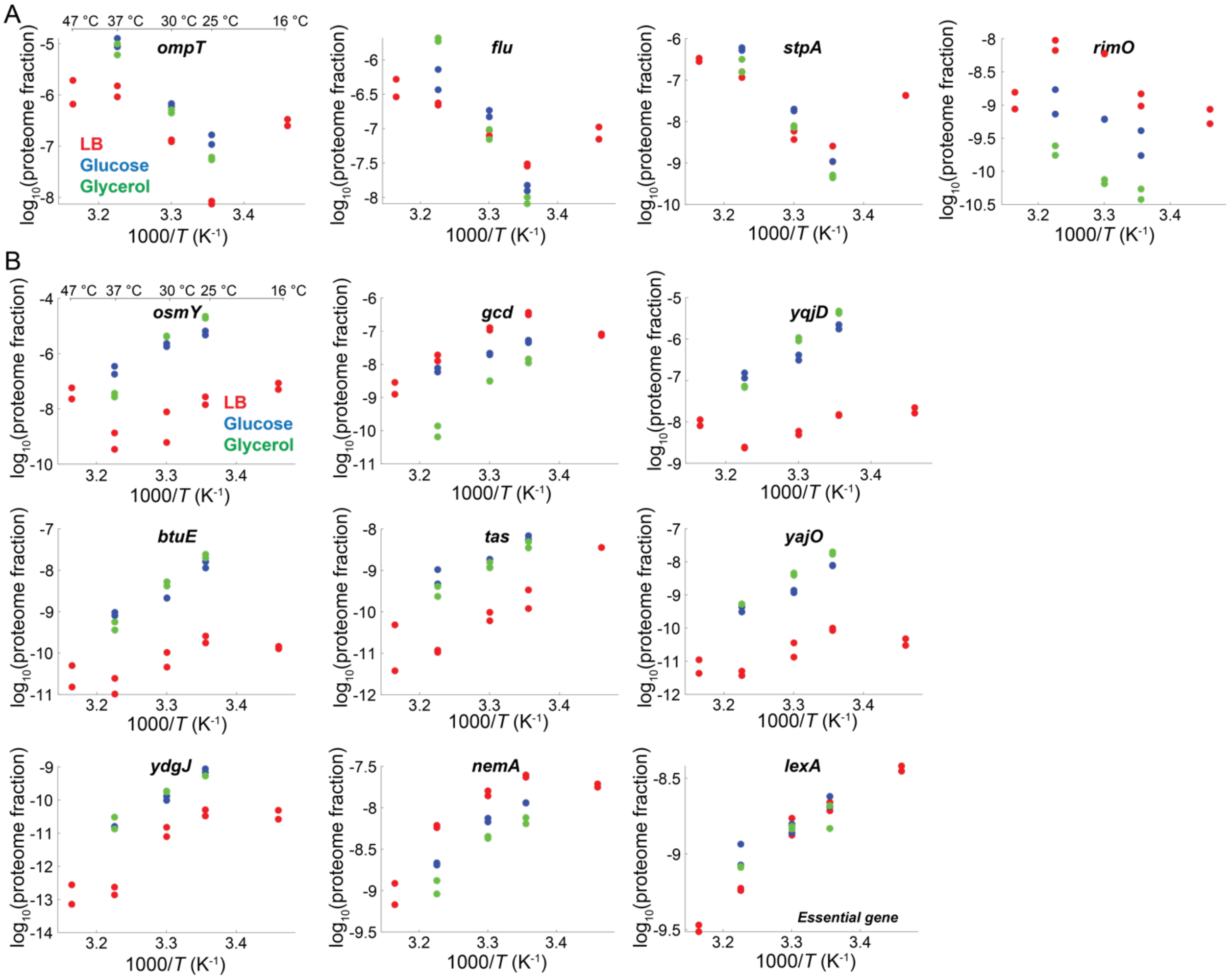


**Figure S9: Proteomics identifies a small subset of proteins whose relative abundance is temperature dependent.**

A) Arrhenius plots (ln(proteome fraction) versus 1/absolute temperature) of the four proteins whose relative abundance increased >2-fold between 25 °C and 37 °C across all media (LB, glucose, glycerol). Temperatures in Celsius are shown for reference in the first plot. Replicates are shown as individual filled circles.

B) Arrhenius plots (ln(proteome fraction) versus 1/absolute temperature) of the nine proteins whose relative abundance decreased >2-fold between 25 °C and 37 °C across all media (LB, glucose, glycerol). Temperatures in Celsius are shown for reference in the first plot. Replicates are shown as individual filled circles.

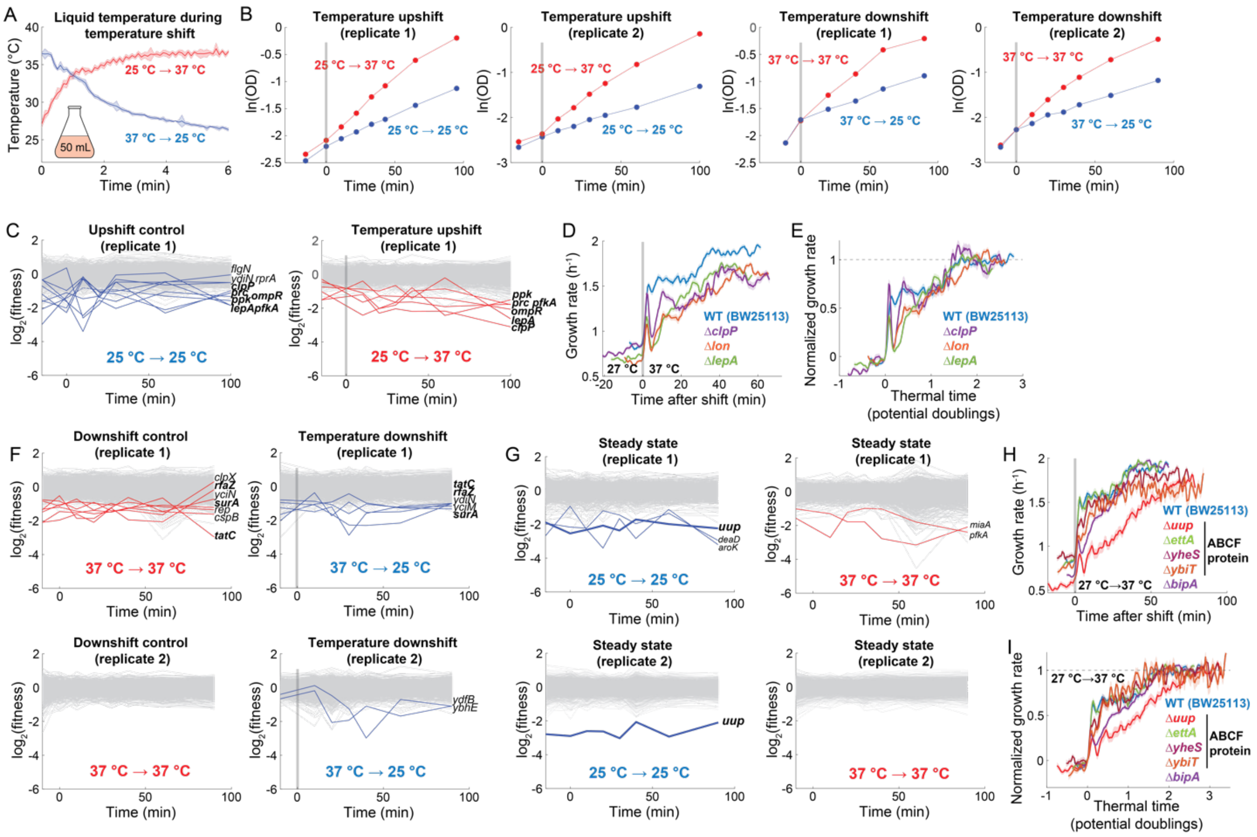


**Figure S10: A pooled transposon library screen did not identify any non-essential genes required for temperature upshift responses.**

A) Temperature readout of 50 mL of water in an Erlenmeyer flask undergoing temperature shifts between 25 °C and 37 °C (Methods). Temperature measurements were averaged over 0.1-s bins and the shaded region represents ±1 standard deviation.

B) Natural logarithm of the optical density of the pooled transposon library during a temperature shift between 25 °C and 37 °C in rich medium (LB), along with a control maintained at the starting temperature. Each time point corresponds to when a sample was collected for sequencing (Methods).

C) Trajectories of $\log_2$(relative abundance) averaged over all mutants in each gene during an upshift from 25 °C to 37 °C. Genes whose average $|\log_2$(relative abundance)| compared with $t$=0 during the upshift experiment was >1 are highlighted in both the control experiment (blue, left) and the upshift (red, right). All genes identified as outliers in the upshift (red, right) were also outliers in the control (blue, left) and are bolded for clarity. Outliers only identified in the control are unbolded. The second biological replicate is shown in Figure 3B,C.

D) Single-cell growth rates of genes identified as the three largest outliers during upshifts across biological replicates from the transposon screen in rich growth medium (LB) during an upshift from 27 °C to 37 °C (Δ*clpP*, purple, *n*=765 cells; Δ*lon*, orange, *n*=1286 cells; Δ*lepA*, green, *n*=1167 cells). The parent BW25113 is shown for comparison (blue, *n*=734 cells). Data are the mean±1 SEM (shaded region) at each time point.

E) Normalized growth rate versus thermal time for each trajectory in (D).

F) Trajectories of $\log_2$(relative abundance) averaged over all mutants in each gene during a downshift from 37 °C to 25 °C. Genes whose average |$\log_2$(relative abundance)| compared with $t$=0 during the downshift experiment was >1 during the experiment are highlighted in both the control (red, left) and the downshift experiment (blue, right). Genes identified as outliers in the downshift (red, right) that were also outliers in the control (right, left) are bolded for clarity. No mutant was present in both biological replicates.

G) Trajectories of $\log_2$(relative abundance) averaged over all mutants in each gene during control experiments. Genes whose average |$\log_2$(relative abundance)| compared with $t$=0 at 37 °C was >1 throughout the experiment is highlighted in color (blue: 25 °C, red: 37 °C). The only gene that was a hit in both biological replicates was *uup*.

H) Single-cell growth rates of knockouts of all ABC-F genes and $\Delta$*bipA* in rich medium (LB) during an upshift from 27 °C to 37 °C. ($\Delta$*uup*, red, *n*=609 cells), ($\Delta$*ettA*, green, *n*=752 cells), ($\Delta$*yheS*, dark red, *n*=797 cells), ($\Delta$*ybiT*, orange, *n*=378 cells), ($\Delta$*bipA*, purple, *n*=1006 cells). The parent BW25113 is shown for comparison (blue, *n*=734 cells). Data are the mean±1 SEM (shaded region) at each time point.

I) Normalized growth rate versus thermal time for each trajectory in (H).

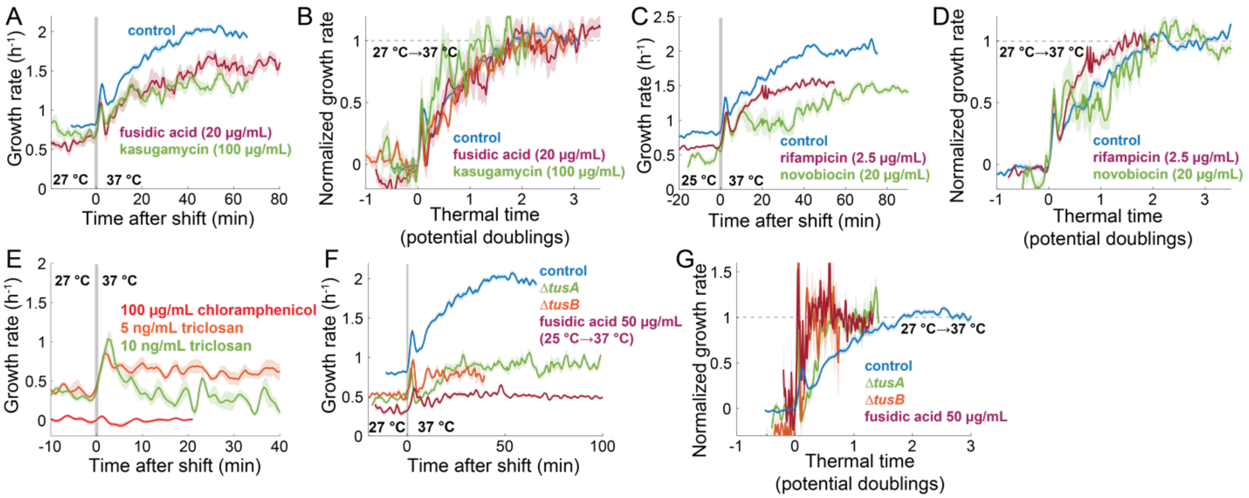


**Figure S11: Normalized upshift response times are largely unaffected by sublethal doses of antibiotics.**

A) Single-cell growth rates of *E. coli* MG1655 on rich medium (LB) with 20 µg/mL fusidic acid (dark red, *n*=132 cells) or 100 µg/mL kasugamycin (green, *n*=79 cells) during an upshift from 27 °C to 37 °C. The untreated control is shown for comparison (blue, *n*=792 cells). Data are the mean±1 SEM (shaded region) at each time point.

B) Normalized growth rate versus thermal time for each trajectory in (A).

C) Single-cell growth rates of *E. coli* MG1655 on rich medium (LB) with 2.5 µg/mL rifampicin (dark red, *n*=522 cells) or 100 µg/mL novobiocin (green, *n*=225 cells) during an upshift from 25 °C to 37 °C. The untreated control is shown for comparison (blue, *n*=773 cells). Data are the mean±1 SEM (shaded region) at each time point.

D) Normalized growth rate versus thermal time for each trajectory in (C).

E) Single-cell growth rates of *E. coli* MG1655 on rich medium (LB) with 5 ng/mL triclosan (orange, *n*=83 cells), 10 ng/mL triclosan (green, *n*=32 cells), or 100 μg/mL chloramphenicol (red, *n*=40 cells) during an upshift from 27 °C to 37 °C. Data are the mean±1 SEM (shaded region) at each time point. We note that at inhibitory concentrations of the ribosome inhibitor chloramphenicol (100 μg/mL), no changes in cell volume occurred during temperature upshifts, indicating that active protein synthesis and/or growth is required for shift dynamics.

F) Single-cell growth rates of wild-type *E. coli* MG1655 on rich medium (LB) with 50 μg/mL fusidic acid (dark red, *n*=86 cells), or BW25113 Δ*tusA* (green, *n*=163 cells) and Δ*tusB* (orange, *n*=118 cells) during an upshift from 27 °C to 37 °C. The untreated control is shown for comparison (blue, *n*=792 cells). Data are the mean±1 SEM (shaded region) at each time point.

G) Normalized growth rate versus thermal time for each trajectory in (F).

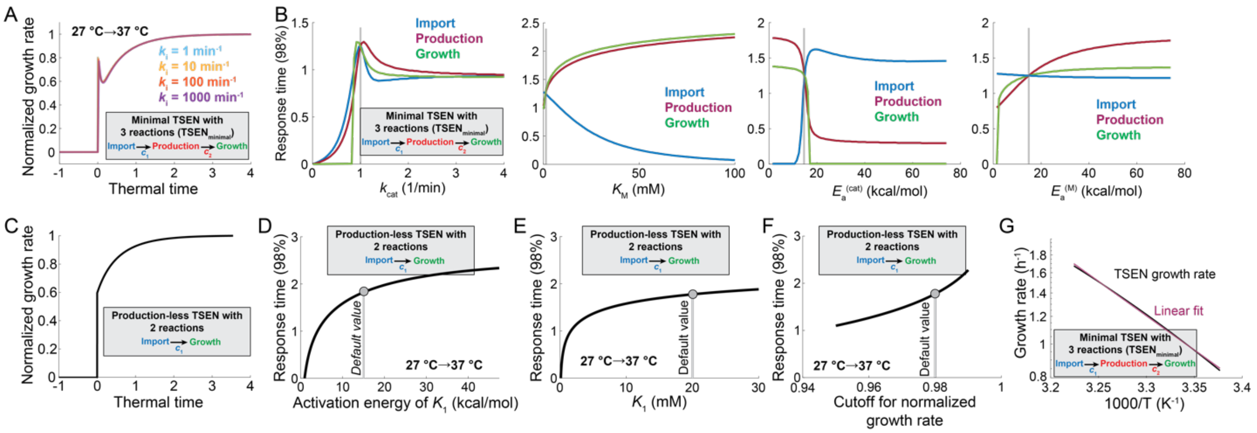


**Figure S12: Effects of changes in TSEN model parameters on temperature-shift response dynamics.**

A) Increasing the catalytic rate ($k_i$) for each reaction in a bottlenecked minimal TSEN model (gray box) from 1 min$^{-1}$ to 1000 min$^{-1}$ has virtually no effect on the response.

B) Effect of model parameters on the normalized response time to a temperature upshift from 27 °C to 37 °C in the minimal TSEN model (gray box) for each reaction (import, production, growth). The definitions of each parameter are provided in Fig. 5A. All other parameters were set to default values (vertical gray bars) in each simulation. Notably, increases in the activation energy of the $K_M$ of the production reaction produced the largest increase in response times across all activation energies (right).

C) The analytically tractable production-less TSEN model (gray box, Supplementary Text) predicts a non-zero response time. The simulation used default parameters ($k_i$= 1 min$^{-1}$, $K_M$ = 1 mM, $E_a^{cat}$ = 15 kcal/mol, $E_a^{M}$= 15 kcal/mol), with the exception of the Michaelis-Menten constant of the second reaction $K_1$ = 20 mM.

D) Normalized response time increases with increased activation energy of $K_1$ in the production-less TSEN model (gray box).

E) Normalized response time increases with increased $K_1$ in the production-less TSEN model (gray box).

F) Normalized response time was between 1 and 2 doublings when the cutoff used to define the adaptation was increased from 95% to 99% of the steady-state growth rate difference in the production-less TSEN model (gray box).

G) Arrhenius plot of steady-state growth rate across temperatures predicted by the minimal TSEN model (gray box) exhibits slightly non-linear behavior.

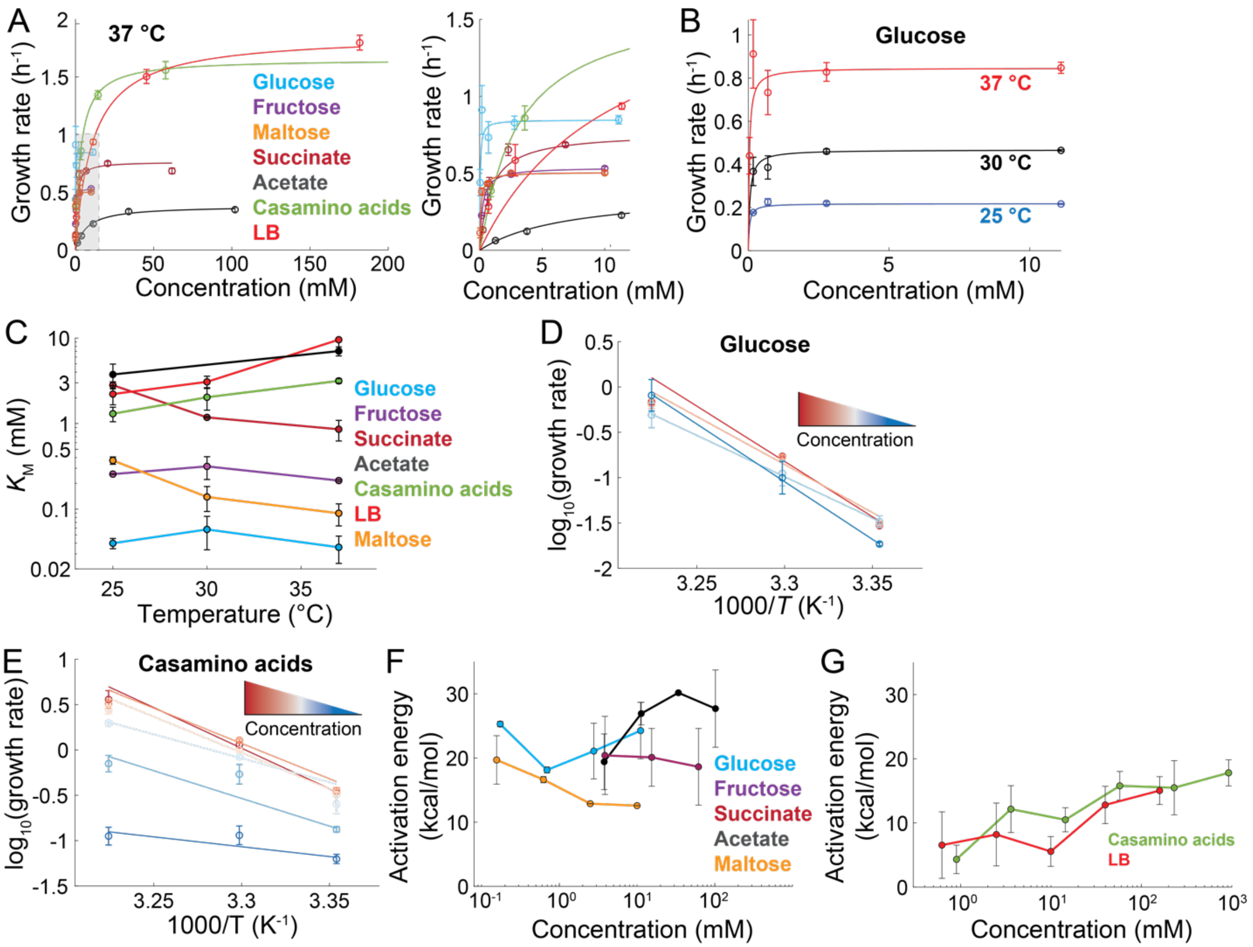


**Figure S13: *E. coli* exhibits Michaelis-Menten kinetics across substrates.**

A) Left: Liquid-culture growth rates of *E. coli* MG1655 grown on a variety of substrates at 37 °C (Methods). Data are the mean of the maximum growth rate extracted from three replicate growth curves and error bars represent ±1 standard deviation (SD). Right: Expanded view of growth rates versus concentration from the outlined box on the left.

B) Liquid-culture growth rates of *E. coli* MG1655 in MOPS minimal medium supplemented with various concentrations of D-glucose at 25 °C (blue), 30 °C (black), or 37 °C (red). Data are the mean of the maximum growth rate extracted from three replicate growth curves and error bars represent ±1 SD.

C) Michaelis-Menten constants ($K_M$) of *E. coli* MG1655 growth rates across growth media and temperatures. Data are estimates from a non-linear weighted fit and error bars represent ±1 SEM.

D) Arrhenius plots of ln(growth rate) versus 1/(absolute temperature) for *E. coli* MG1655 grown in MOPS minimal medium supplemented with glucose at concentrations between 0.17 mM and 11 mM (blue-to-red). Data are the mean across three replicates and error bars represent ±1 SD. Weighted linear fits were performed for each concentration.

E) Arrhenius plots of ln(growth rate) versus 1/(absolute temperature) for *E. coli* MG1655 grown in MOPS minimal medium supplemented with casamino acids at concentrations between 0.9 mM and 923 mM (blue-to-red). Data are the mean across three replicates and error bars represent ±1 SD. Weighted linear fits were performed for each concentration.

F) Activation energy versus substrate concentration of *E. coli* MG1655 grown in minimal media without amino acids supplemented with glucose, fructose, acetate, succinate, or maltose. Activation energies are estimates from a linear weighted fit of Arrhenius plots and error bars represent ±1 SEM.

G) Activation energy versus substrate concentration of *E. coli* grown in LB (red) or MOPS minimal medium supplemented with casamino acids (green). Activation energies are estimates from a linear weighted fit of Arrhenius plots and error bars represent ±1 SEM.

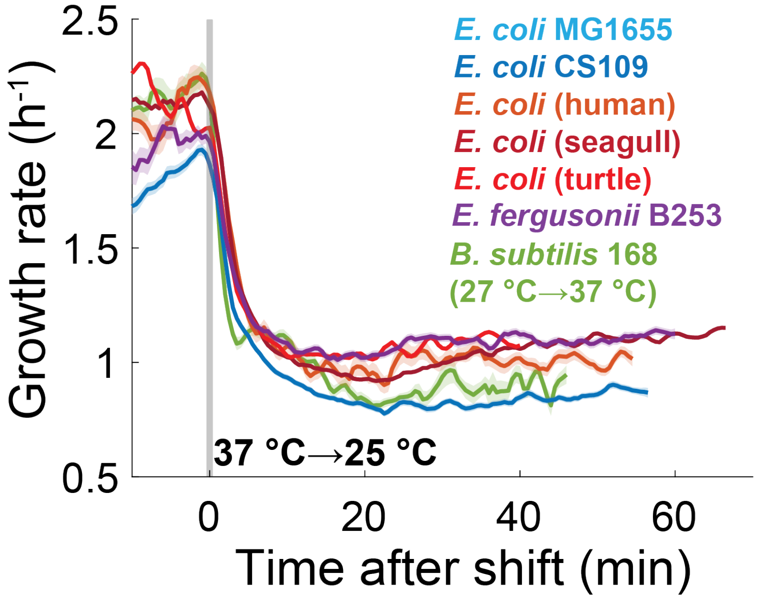


**Figure S14: Growth rate response to a temperature downshift is rapid across organisms.**

Single-cell growth rate response to a temperature downshift on rich medium (LB) of laboratory-evolved (blue, CS109, *n*=911 cells) and naturally isolated *E. coli* strains (orange to red, *n*=417-1186 cells), *Escherichia fergusonii* (purple, *n*=902 cells), and *Bacillus subtilis* (green, *n*=117 cells). All downshifts were from 37 °C to 25 °C, except for *B. subtilis* (37 °C to 27 °C). Data are the mean±1 SEM (shaded region) at each time point.

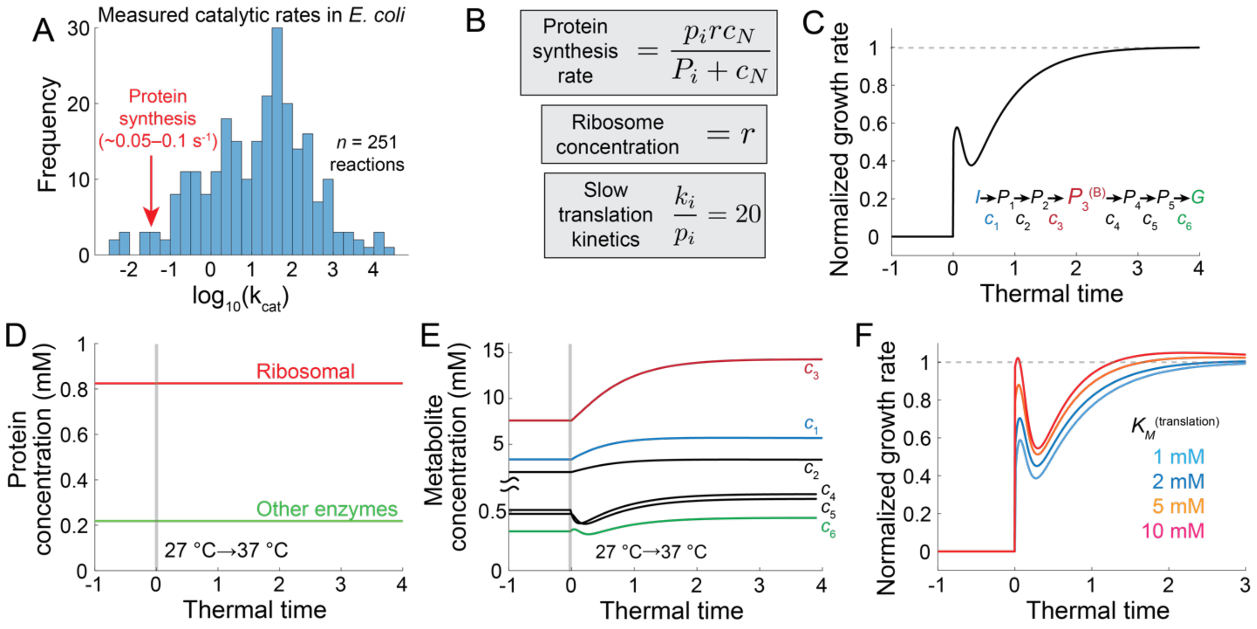


**Figure S15: A TSEN model with protein synthesis predicts a temperature-invariant proteome.**

A) Measured catalytic rates for 251 wild-type enzymes from *E. coli* (BRENDA database) (Supplementary Text). Labeled in red is the estimated protein synthesis rate for a single ribosome.

B) Protein synthesis follows Michaelis-Menten kinetics in the TSEN model. Translation of new protein (index $i$) by ribosomes ($r$) is defined by the catalytic rate (i.e., translational elongation), $p_i$, and Michaelis-Menten constant, $P_i$. The catalytic rate of the ribosome is constrained as 20-fold less than other enzymatic catalytic rates (Supplementary Text).

C) Normalized growth rate versus thermal time for a simulation of a full bottlenecked TSEN model similar to Fig. 5C with translation. Parameter values for translation are described in the Supplementary Text, and all other parameters were set to the same values as in Fig. 5C.

D) Protein concentration remained virtually constant after a temperature upshift in the simulation of the full bottlenecked TSEN with translation in (C).

E) Metabolite concentrations after a temperature upshift in the simulation of the full bottlenecked TSEN with translation in (C) exhibited similar dynamics to the model with constant enzyme concentrations in Fig. 5E.

F) Increasing the Michaelis-Menten constant ($P_i$) for translation decreased the response time and increased the normalized spike height. All other parameters were set to the same values as in (C).

### SUPPLEMENTARY TEXT

#### ***Enzymatic reactions in the Temperature-Sensitive Enzyme Network (TSEN)*** **are** ***assumed to follow the Arrhenius equation***

The Arrhenius equation is an empirical model from equilibrium thermodynamics[4] that describes the rate of a reaction ($k$) as a function of temperature ($T$) according to

$$k(T) = k_o e^{-\frac{E_a}{k_B T}} \tag{1}$$

where $k_o$ is a constant, $E_a$ is the activation energy for the reaction, and $k_B$ is the Boltzmann constant. Increases in $E_a$ or decreases in $T$ decrease the reaction rate, such that $E_a$ describes the temperature sensitivity of the reaction. This empirical law can be derived from transition-state theory, which has an analogous formulation known as the Eyring equation[5].

The vast majority of reactions have catalytic rates that obey the Arrhenius equation[6], and diverse organisms typically have a temperature range over which their growth rate obeys the Arrhenius equation[4,7]. Thus, in our model we assumed that the temperature dependence of all reactions follows the Arrhenius equation (Eq. 1).

#### ***Each reaction obeys Michaelis-Menten kinetics***

Most biological enzymes obey Michaelis-Menten (MM) kinetics[8]. As such, MM kinetics are widely used as the default framework for reporting enzyme kinetic parameters[9]. Thus, in our model we assumed that each reaction obeys MM kinetics and assayed the effects of the reaction being above or below saturation.

In our enzyme network, each intermediate metabolite, represented by the concentration $c_{i+1}$, is produced by the $i^{\text{th}}$ reaction according to

$$c_i + e_i \rightleftharpoons (c_i e_i)^* \longrightarrow c_{i+1} + e_i \qquad (2)$$

where $e_i$ free enzymes bind $c_i$ to form an intermediate state $(c_i e_i)^*$ with a forward rate $f_i$ and reverse rate $r_i$. The intermediate state then leads to production of $c_{i+1}$ at the catalytic rate $k_i$. The production rate of $c_{i+1}$ can be derived by assuming that the concentration of the intermediate state $(c_i e_i)^*$ is approximately constant[10], which requires

$$\frac{d(c_i e_i)^*}{dt} = f_i c_i e_i - r_i (c_i e_i)^* - k_i (c_i e_i)^* = 0.$$

Solving for the intermediate state,

$$(c_i e_i)^* = \frac{E_i c_i}{\frac{r_i + k_i}{f_i} + c_i} \qquad (3)$$

where $E_i = e_i + (c_i e_i)^*$ is the total enzyme concentration. Since we determined that the proteome is largely constant across Arrhenius temperatures (Fig. 2), we initially assume that enzyme concentrations are constant in our TSEN model; note that we show in a later section of this text that this assumption is also a natural consequence of our model. The production rate of $c_{i+1}$ is the well-known Michaelis-Menten equation,

$$\frac{dc_{i+1}}{dt} = k_i (c_i e_i)^* = \frac{k_i E_i c_i}{K_i + c_i} \qquad (4)$$

where $K_i = \frac{r_i + k_i}{f_i}$ is the MM constant of the reaction.

#### *Chained MM reactions obey simple dynamics*

In chained reactions, a substrate $c_i$ is produced by the ($i$-1)$^{th}$ reaction according to Eq. 2 and then consumed by a subsequent reaction:

$$c_{i-1} + e_{i-1} \rightleftharpoons (c_{i-1}e_{i-1})^* \longrightarrow c_i + e_{i-1}$$

$$c_i + e_i \rightleftharpoons (c_i e_i)^* \longrightarrow c_{i+1} + e_i.$$

We assume that each $(c_i e_i)^*$ obeys the steady-state approximation in Eq. 3 to obtain

$$\frac{dc_i}{dt} = k_{i-1}(c_{i-1}e_{i-1})^* + r_i(c_i e_i)^* - f_i c_i e_i.$$

The last two terms are equal to $k_i(c_i e_i)^*$, as shown above. Therefore,

$$\frac{dc_i}{dt} = k_{i-1}(c_{i-1}e_{i-1})^* - k_i(c_i e_i)^*.$$

Again applying Eq. 3 for each $(c_i e_i)^*$,

$$\frac{dc_i}{dt} = \frac{k_{i-1}E_{i-1}c_{i-1}}{K_{i-1} + c_{i-1}} - \frac{k_i E_i c_i}{K_i + c_i}. \quad (5)$$

Eq. 5 describes the dynamics of intermediate substrates produced and consumed by enzymes at a fixed concentration within a fixed volume. The first term on the right-hand side describes production by enzyme $E_{i-1}$ and the second term describes consumption by enzyme $E_i$[11].

***Import and growth reactions obey MM kinetics***

Many transporters are known to obey MM kinetics[9]. For example, the *E. coli* maltose transporter complex consumes ATP with a $K_M$~300 μM, whose catalytic rate is 468 min$^{-1}$ at 37 °C and exhibits Arrhenius behavior with $E_a$~14 kcal/mol[12]. The glucose-specific transporter in *E. coli*, PtsG, has a $K_M$~60 μM[13], in agreement with our measurement of $K_M$ for growth on glucose (~30 μM) (Fig. S13C). Thus, we assume the import reaction in our model consumes external metabolites $c_0$ and produces the first intermediate metabolite $c_1$ at a rate

$$\frac{dc_1}{dt} = \frac{k_0 E_0 c_0}{K_0 + c_0}. \tag{6}$$

We assume that the external metabolite pool is sufficiently large that $c_0$ is constant during steady-state growth, which is reasonable for the agarose hydrogel environment in our single-cell experiments.

Growth reactions also likely follow MM kinetics. For example, the glycosyltransferase PBP1b in *E. coli*, a transmembrane protein that catalyzes polymerization of peptidoglycan, obeys MM kinetics with a $K_{\mathrm{M}}$~20 μM[14]. In our model, the last reaction in the network produces cell volume expansion through incorporation of the final metabolite $c_N$ into cell envelope material, which results in the growth rate

$$g = \gamma_0 \frac{k_N E_N c_N}{K_N + c_N} \tag{7}$$

where $\gamma_0$ is a growth efficiency factor, which we estimate below to be ~0.03 mM$^{-1}$ for *E. coli*.

***Intracellular metabolites are diluted by growth***

While Eq. 5 describes metabolite production and consumption under constant volume conditions, cell volume increases exponentially during steady-state growth[15]. For a single metabolite, its absolute number $N_i$ increases within increasing volume $V$, such that concentration $c_i = \frac{N_i}{V}$ and

$$\frac{dc_i}{dt} = \frac{1}{V}\frac{dN_i}{dt} - \frac{N_i}{V}\frac{1}{V}\frac{dV}{dt}.$$

The first term on the left-hand side is the constant-volume production rate of $c_i$ and the second term is the product of the instantaneous concentration $\frac{N_i}{V}$ and the instantaneous growth rate $g = \frac{1}{V}\frac{dV}{dt}$:

$$\frac{dc_i}{dt} = \frac{dc_i}{dt}\Big|_{\text{constant } V} - c_i g = \frac{k_{i-1}E_{i-1}c_{i-1}}{K_{i-1} + c_{i-1}} - \frac{k_i E_i c_i}{K_i + c_i} - c_i g. \quad (8)$$

***Steady-state growth rate depends only on import and the total concentration of intracellular metabolites***

At steady state, all metabolite concentrations are constant $\left(\frac{dc_i}{dt} = 0 \text{ for all } i\right)$, hence

$$\sum_{i=1}^{N} \frac{dc_i}{dt} = 0,$$

which can be solved by defining $\alpha_i = \frac{k_i E_i c_i}{K_i + c_i}$:

$$\sum_{i=1}^{N} \frac{dc_i}{dt} = \sum_{i=1}^{N} (\alpha_{i-1} - \alpha_i - g c_i) = \alpha_0 - \alpha_N - g \sum_{i=1}^{N} c_i = 0.$$

Thus, $\alpha_0 - \alpha_N = g \sum_{i=1}^{N} c_i$ and $g = \gamma_0 \alpha_N$, hence $\alpha_0 = \frac{g}{\gamma_0} + g \sum_{i=1}^{N} c_i$ and

$$g = \frac{\alpha_0}{\frac{1}{\gamma_0} + \sum_{i=1}^{N} c_i}. \quad (9)$$

Therefore, the growth rate only depends on import ($\alpha_0$) and the total concentration of intracellular metabolites ($\sum_{i=1}^{N} c_i$), indicating that growth comes at the cost of producing and storing all intermediate metabolites.

***Estimating growth rate from experimental measurements of metabolite uptake***

To test the validity of Eq. 9, we estimated the growth rate of *E. coli* grown on glucose as its sole source of carbon. Importantly, glucose uptake rate is linearly correlated with growth rate in *E. coli*[16], indicating that our model captures the dependence of growth rate on import. The catalytic rate of the major *E. coli* sugar transporter PtsI has been measured at ~12,000 min$^{-1}$ (ref. [17]), consistent with estimates of the glucose uptake rate in *E. coli* at 37 °C (~2000-3600 mM/h)[16,18,19]. The glucose uptake rate of *E. coli* at 30 °C has been estimated at 23 nmol/min/mg dry weight, which is equivalent to ~420 mM/h for a single cell[13]. However, single-cell measurements of *E. coli* grown at 30 °C placed the glucose uptake rate at ~120 mM/h[20]. Thus, the rate of glucose import is likely between 120 mM/h and 3600 mM/h. Based on our estimate of $\gamma_0 \leq 0.02$ mM$^{-1}$ and estimates of the total metabolite concentration in *E. coli* during growth on glucose (~300 mM)[21], Eq. 9 predicts a growth rate between ~0.3–10 h$^{-1}$, roughly consistent with our single-cell and liquid-culture measurements of 0.8–1 h$^{-1}$ at 37 °C (Fig. 1G, S13A,B). Alternatively, for an import rate of 420–3600 mM/h, a growth rate of 0.8 h$^{-1}$ implies that $\gamma_0$ is 0.0002-0.006 mM$^{-1}$.

We can also estimate the glucose uptake rate during glucose-limited growth from the elemental composition of *E. coli*, which is ~40% carbon by mass[22]. During early log phase growth on glucose, cellular biomass is ~150 fg, which corresponds to ~60 fg of carbon[22]. Since cells have a volume of ~1 $\mu$m$^3$, the carbon molarity is ~5 M. *E. coli* growth rate on glucose at 37 °C is 0.6–1.0 h$^{-1}$, corresponding to a carbon growth rate of 3–5 M/h, which must be matched by carbon import to achieve balanced growth. Since each glucose has 6 carbons, the glucose import rate must be at least 500–800 mM/h, largely in agreement with the measured range from the literature. By constraining $\gamma_0$ by our measured liquid-culture growth rate on glucose (0.8 h$^{-1}$, Fig. S13A,B), an import rate of 500 mM/h gives an estimate $\gamma_0$~0.004 mM$^{-1}$. In our model, the import rate at saturation is determined by $k_i E_i$, equal to 60 mM/h under default parameters. Thus, $\gamma_0$ for the minimal TSEN can be scaled by physiological values for the import rate (~500 mM/h), such that

$$\gamma_0^{\mathrm{model}} \approx \left(\frac{500}{60}\right) 0.004\ \mathrm{mM}^{-1} = 0.03\ \mathrm{mM}^{-1}.$$

***Network branching does not affect growth rate in the TSEN model***

We also considered a TSEN model with a single branch point, wherein a component $c_p$ is consumed by two enzymes ($e_{a,0}$, $e_{b,0}$), which lead to separate branches (*a,b*) of lengths *L* and *M*, respectively. This structure is representative of e.g., parallel metabolism between glycolysis and the pentose phosphate pathway[23]. For simplicity, we assume that each branch eventually contributes separately to growth through a final reaction similar to Eq. 7.

The concentration of $c_p$ varies in time according to

$$\frac{dc_p}{dt} = \alpha_{p-1} - \alpha_{a,0} - \alpha_{b,0} - c_p g,$$

where each $\alpha$ is a parametrization of MM kinetics used in deriving Eq. 9. Each branch (*a,b*) results in growth according to

$$g_a = \gamma_a \alpha_{a,L} = \gamma_a \frac{k_{a,L} E_{a,L} c_{a,L}}{K_{a,L} + c_{a,L}} \text{ and } g_b = \gamma_b \alpha_{b,M} = \gamma_b \frac{k_{b,M} E_{b,M} c_{b,M}}{K_{b,M} + c_{b,M}},$$

such that $g = g_a + g_b$. At steady state, all concentrations are constant $\left(\sum_{i=1}^{N} \frac{dc_i}{dt} = 0\right)$, such that

$$\sum_{i=1}^{p-1}(\alpha_{i-1} - \alpha_i - gc_i) + \left(\alpha_{p-1} - \alpha_{a,0} - \alpha_{b,0} - c_p g\right) + \sum_{i=1}^{L}(\alpha_{a,i-1} - \alpha_{a,i} - gc_{a,i}) +$$

$$\sum_{i=1}^{M}(\alpha_{b,i-1} - \alpha_{b,i} - gc_{b,i}) = 0,$$

hence

$$\alpha_0 - \alpha_{a,L} - \alpha_{b,M} - g\sum_{i=1}^{p} c_i - g\sum_{i=1}^{L} c_{a,i} - g\sum_{i=1}^{M} c_{b,i} = 0.$$

Therefore,

$$\alpha_0 - \frac{g_a}{\gamma_a} - \frac{g_b}{\gamma_b} - g\sum_{j}^{p+L+M} c_j = 0.$$

Assuming that each growth reaction has the same efficiency factor ($\gamma_0 = \gamma_a = \gamma_b$),

$$g_{\text{branched}} = \frac{\alpha_0}{\frac{1}{\gamma_0} + \sum_{j}^{p+L+M} c_j}. \qquad (10)$$

This equation is equivalent to Eq. 9, except that the sum includes all metabolites in each branch. Thus, branches decrease the steady-state growth rate, dependent on the length and kinetic properties of the branches.

Note that Eq. 10 can be obtained similarly for an arbitrary number of branch points that lead to growth with the same growth efficiency factor. Furthermore, if branches all funnel to a final metabolite that is consumed by a final growth reaction, Eq. 10 still holds.

***Production-less TSEN model***

Consider a model of cell growth involving only import and expansion reactions (i.e., without the potential for spike behavior), such that there is a single intermediate metabolite with concentration $c_1$ whose dynamics are governed by the equation

$$\frac{dc_1}{dt} = I_0 - \frac{k_1 e_1 c_1}{K_1 + c_1} - c_1 g, \tag{11}$$

where $I_0$ is the import rate and $g$ is the growth rate

$$g = \gamma_0 \frac{k_1 e_1 c_1}{K_1 + c_1}. \tag{12}$$

Substitution of Eq. 12 into Eq. 11 and solving for *dt* gives

$$dt = \frac{dc_1}{I_0} \frac{(K_1 + c_1)}{K_1 + c_1\left(1 - \frac{k_1 e_1}{I_0}\right) - \frac{\gamma_0 k_1 e_1}{I_0} c_1^2}. \tag{13}$$

*Well-balanced approximation*

If we assume that the import rate $I_0$ and the saturated consumption rate $k_1 e_1$ are approximately equal (i.e., well-balanced), i.e.,

$$\frac{k_1 e_1}{I_0} = 1, \tag{14}$$

then the steady-state concentration at temperature $T$ is

$$c_1^{\mathrm{ss}}(T) = \sqrt{\frac{K_1(T)}{\gamma_0}}. \tag{15}$$

In this scenario, Eq. 13 becomes

$$dt = \frac{dc_1}{I_0} \frac{(K_1 + c_1)}{K_1 - \gamma_0 c_1^2}. \tag{16}$$

The growth rate at steady state is

$$g^{\mathrm{ss}} = \frac{I_0}{\frac{1}{\gamma_0} + c_1^{\mathrm{ss}}} \tag{17}$$

and the doubling time is

$$\tau_D^{\mathrm{ss}} = \frac{\frac{1}{\gamma_0} + c_1^{\mathrm{ss}}}{I_0} \ln 2\,. \tag{18}$$

*The response time under the well-balanced approximation*

To define the response time, we solve for the time $t$ required to reach a threshold of $c_1^{\mathrm{ss}}(T_2)$, defined as $\overline{c_1}(T_2)$, starting from steady state at the lower temperature, $c_1^{\mathrm{ss}}(T_1)$, assuming that all kinetics after the switch are set by $T_2$. This response time can be obtained by integrating Eq. 16:

$$t = \int_{c_1^{\mathrm{ss}}(T_1)}^{\overline{c_1}(T_2)} \frac{dc_1}{I_0} \frac{(K_1 + c_1)}{K_1 - \gamma_0 c_1^2} \tag{19}$$

and using Eq. 15,

$$\int \frac{dc_1}{I_0} \frac{(K_1 + c_1)}{K_1 - \gamma_0 {c_1}^2} = \frac{1}{I_0}\sqrt{\frac{K_1}{\gamma_0}} \tanh^{-1} \frac{c_1}{\sqrt{K_1/\gamma_0}} - \frac{1}{2\gamma_0 I_0} \ln(K_1 - \gamma_0 c_1^2).$$

Using Eq. 15, and the assumption that kinetics are evaluated at $T_2$,

$$\int \frac{dc_1}{I_0} \frac{(K_1 + c_1)}{K_1 - \gamma_0 {c_1}^2} = \frac{1}{I_0} c_1^{\mathrm{ss}}(T_2) \tanh^{-1} \frac{c_1}{c_1^{\mathrm{ss}}(T_2)} - \frac{1}{2\gamma_0 I_0} \ln(K_1 - \gamma_0 c_1^2).$$

Assuming that $\overline{c_1}(T_2) < c_1^{\mathrm{ss}}(T_2)$,

$$\tanh^{-1} \frac{c_1}{c_1^{\mathrm{ss}}(T_2)} = \frac{1}{2} \ln\left(\frac{c_1^{\mathrm{ss}}(T_2) + c_1}{c_1^{\mathrm{ss}}(T_2) - c_1}\right).$$

Hence,

$$\int \frac{dc_1}{I_0} \frac{(K_1 + c_1)}{K_1 - \gamma_0 {c_1}^2} = c_1^{\mathrm{ss}}(T_2) \frac{1}{2I_0} \ln\left(\frac{c_1^{\mathrm{ss}}(T_2) + c_1}{c_1^{\mathrm{ss}}(T_2) - c_1}\right) - \frac{1}{2\gamma_0 I_0} \ln(K_1 - \gamma_0 c_1^2)$$

which we evaluate at the integral’s limits to obtain

$$\int_{c_1^{ss}(T_1)}^{\overline{c_1}(T_2)} \frac{1}{I_0} \frac{dc_1(K_1 + c_1)}{K_1 - \gamma_0 c_1^2}$$

$$= \frac{1}{2\gamma_0 I_0}\left(c_1^{ss}(T_2)\gamma_0 \ln\left(\frac{c_1^{ss}(T_2) + \overline{c_1}(T_2)}{c_1^{ss}(T_2) - \overline{c_1}(T_2)} \cdot \frac{c_1^{ss}(T_2) - c_1^{ss}(T_1)}{c_1^{ss}(T_2) + c_1^{ss}(T_1)}\right)\right.$$

$$\left. - \ln\left(\frac{c_1^{ss}(T_2)^2 - \overline{c_1}^2(T_2)}{c_1^{ss}(T_2)^2 - c_1^{ss}(T_1)^2}\right)\right).$$

We assume that $K_1(T)$ is an Arrhenius function with activation energy $E_1$:

$$K_1(T) = K^* e^{-\frac{E_1}{k_B T}}$$

and the steady-state concentration of $c_1^{ss}(T_1)$ can be related to $c_1^{ss}(T_2)$ using Eq. 5 as

$$\frac{c_1^{ss}(T_1)}{c_1^{ss}(T_2)} = e^{-\frac{E_a}{2k_B\left(\frac{1}{T_1} - \frac{1}{T_2}\right)}} \equiv f_1 \,. \qquad (20)$$

Finally, we define the threshold $\overline{c_1}(T_2)$ (Eq. 19) to be between $c_1^{ss}(T_1)$ and $c_1^{ss}(T_2)$, such that

$$\overline{c_1}(T_2) = f c_1^{ss}(T_2).$$

Now the time $t$ to reach $\overline{c_1}(T_2)$ is

$$t = \frac{1}{2\gamma_0 I_0}\left(c_1^{ss}\gamma_0 \ln\left(\frac{1+f}{1-f} \cdot \frac{1-f_1}{1+f_1}\right) - \ln\frac{1-f^2}{1-f_1^2}\right). \qquad (21)$$

Defining the response time ($\tau_R$) as the time for $c_1(t, T_2)$ to reach $\overline{c_1}(T_2)$ normalized by the steady-state doubling time at $T_2$ (Eq. 18),

$$\tau_R = \frac{1}{(1 + \gamma_0 c_1^{ss}) 2\ln 2}\left(c_1^{ss}\gamma_0 \ln\left(\frac{1+f}{1-f} \cdot \frac{1-f_1}{1+f_1}\right) - \ln\frac{1-f^2}{1-f_1^2}\right).$$

Alternatively,

$$\tau_R = \frac{1}{\left(1+\sqrt{\gamma_0 K_1(T_2)}\right)2\ln 2}\left(\sqrt{\gamma_0 K_1(T_2)}\ln\left(\frac{1+f}{1-f}\cdot\frac{1-f_1}{1+f_1}\right)-\ln\frac{1-f^2}{1-f_1^2}\right). \qquad (22)$$

*Predictions of model*

To determine the effects of each parameter ($\gamma_0, K_1(T_2), f$) on the response time ($\tau_R$), we evaluated Eq. 22 across a range of values. As in the full TSEN model, we set the default activation energy to 15 kcal/mol to match physiological conditions. Similarly, we set $\gamma_0 = 0.02$ mM$^{-1}$ to match the full TSEN model, and $K_1(T_2)$=20 mM to match the bottleneck reaction in the full TSEN model.

To examine the response time ($\tau_R$) as a function of the growth rate as it nears steady state at $T_2$, we define a threshold of the normalized growth rate ($f_g$), defined as

$$f_g \equiv \frac{g\left(\overline{c_1}(T_2)\right)-g\left(c_1(T_1)\right)}{g\left(c_1(T_2)\right)-g\left(c_1(T_1)\right)}. \qquad (23)$$

Combining Eq. 17 with Eq. 23 and using the definition of $\overline{c_1}(T_2) = f c_1^{\mathrm{ss}}(T_2)$ yields

$$f = \frac{1}{\gamma_0 c_1^{\mathrm{ss}}(T_1)}\left(\frac{\frac{I_0(T_2)}{I_0(T_1)}\left(1+\gamma_0 c_1^{\mathrm{ss}}(T_1)\right)}{1+f_g\left(\frac{I_0(T_2)}{I_0(T_1)}\frac{1+\gamma_0 c_1^{\mathrm{ss}}(T_1)}{1+\gamma_0 c_1^{\mathrm{ss}}(T_2)}-1\right)}-1\right),$$

where $I_0(T)$ is the import rate at temperature $T$. Assuming that the import rate follows Arrhenius behavior with an activation energy $E_a$,

$$\mu \equiv \frac{I_0(T_2)}{I_0(T_1)} = e^{-\frac{E_a}{k_B}\left(\frac{1}{T_2}-\frac{1}{T_1}\right)}$$

and using Eq. 15 for the steady-state concentration,

$$f = \frac{1}{\sqrt{\gamma_0 K_1(T_2)}}\left(\frac{\mu\left(1+\sqrt{\gamma_0 K_1(T_1)}\right)}{1+f_g\left(\mu\frac{1+\sqrt{\gamma_0 K_1(T_1)}}{1+\sqrt{\gamma_0 K_1(T_2)}}-1\right)}-1\right). \tag{24}$$

Eq. 24 then allows us to evaluate Eq. 22 as a function of the threshold for the normalized growth rate to reach its new steady state.

To determine the effect of the choice of $f_g$, we evaluated the response time for $f_g$ between 0.95 and 0.99. The response time varied between 1.1 and 2.3 doublings (Fig. S12F), which is in reasonable agreement with our measurements (~1.5 doublings) (Fig. 1F,H).

This two-step production-less TSEN model accurately predicts a normalized response time that does not depend on the growth rate at the final temperature (Eq. 22) and produces response times between 1–2 doublings across a large range of parameter values (Fig. S12D–F). We can understand why response times lie between 1–2 doublings by solving for $g(c_1(t))$ from Eq. 17 and 21 as follows:

$$2\gamma_0 I_0 t = \gamma_0 c_1^{\mathrm{ss}}(T_2)\ln\left(\frac{c_1^{\mathrm{ss}}(T_2)-c_1^{\mathrm{ss}}(T_1)}{c_1^{\mathrm{ss}}(T_2)-c_1}\right)$$

$$+\gamma_0 c_1^{\mathrm{ss}}(T_2)\ln\left(\frac{c_1^{\mathrm{ss}}(T_2)+c_1}{c_1^{\mathrm{ss}}(T_2)+c_1^{\mathrm{ss}}(T_1)}\right)+\ln\left(\frac{c_1^{\mathrm{ss}}(T_2)^2-c_1^{\mathrm{ss}}(T_1)^2}{c_1^{\mathrm{ss}}(T_2)^2-c_1^2}\right).$$

The first term on the right-hand side dominates as $c_1$ approaches $c_1^{ss}(T_2)$, hence

$$2\gamma_0 I_0 t \approx \gamma_0 c_1^{ss}(T_2)\ln\left(\frac{c_1^{ss}(T_2)-c_1^{ss}(T_1)}{c_1^{ss}(T_2)-c_1}\right).$$

Solving for $c_1(t, T_2)$,

$$c_1(t,T_2) \approx c_1^{ss}(T_2) - \left(c_1^{ss}(T_2) - c_1^{ss}(T_1)\right)e^{-\frac{2I_0}{\gamma_0}t}. \quad (25)$$

Using the definition of the doubling time from Eq. 17, the decay constant becomes

$$\frac{2I_0}{\gamma_0} = \frac{2\ln 2}{\tau_D}\left(1 + \frac{1}{\gamma_0 c_1^{ss}(T_2)}\right). \quad (26)$$

Eq. 26 indicates that the growth rate, which follows

$$g\left(c_1(t,T_2)\right) = \frac{I_0}{\frac{1}{\gamma_0} + c_1(t,T_2)},$$

also has the same decay constant, such that the growth rate approaches its new steady state exponentially with a decay constant according to Eq. 26. If $\gamma_0 c_1^{ss}(T_2) \gg 1$, then the decay constant is

$$\frac{2I_0}{\gamma_0} \approx \frac{2\ln 2}{\tau_D} = \frac{1.4}{\tau_D},$$

hence $c_1(t,T_2)$ reaches 99% of its steady-state value $c_1^{ss}(T_2)$ within ~3.3 doublings. This value is independent of any choice of kinetic values and potentially sets an upper limit for observed response times in the growth rate. Furthermore, for physiological values based on *E. coli* growth in glucose[21], $\gamma_0 = 0.0002—0.006$ mM$^{-1}$ and $c_{\text{total}} = 300$ mM produces an estimate of 0.5–1.4 doublings for total metabolite concentration to reach 99% of its steady-state value, in reasonable agreement with our measurements of growth-rate response (Fig. 1F,H).

Importantly, since $c_1$ depends only on $K_1$ at steady state (Eq. 15), the production-less TSEN predicts that the temperature sensitivity of the metabolic network is determined by the temperature sensitivity of the Michaelis-Menten constant (Fig. S12C, 7F).

Furthermore, the normalized response time only depends weakly on the $E_a$ of $K_1$ above ~20 kcal/mol and $K_1$>10 mM (Fig. S12C–E).

***TSEN model produces the same dynamics with the inclusion of translation***

While we have so far imposed that enzyme concentrations are constant during a temperature shift, we also considered a TSEN that includes translation dynamics. We modified the model by adding a simple translation mechanism to account for enzyme production (Fig. S15). We assume that binding of ribosomes to mRNA follows MM kinetics, which has been demonstrated experimentally with an estimated $K_{\text{M}}$~3–8.5 μM[24,25].

For each enzyme ($e_i$), its synthesis rate follows MM kinetics with a catalytic rate $\rho_i$ and MM constant $P_i$. This reaction is catalyzed by a fraction, $f_i$, of the total ribosome concentration, $r$. For simplicity, we assume that the final product of the network, $c_N$, is consumed by either the growth reaction (Eq. 7) or by translation:

$$\frac{de_i}{dt} = \frac{p_i(f_i r)c_N}{P_i + c_N}. \tag{27}$$

We also assume that ribosome production is limited by ribosomal protein synthesis, such that it follows MM kinetics with catalytic rate $p_R$ and MM constant $P_R$, and is produced by a fraction, $f_R$, of the total ribosome concentration.

$$\frac{dr}{dt} = \frac{p_R(f_R r)c_N}{P_R + c_N}. \tag{28}$$

Each enzyme, including the ribosome, is diluted by growth such that its dynamics are determined by

$$\frac{de_i}{dt} = \frac{p_i(f_i r)c_N}{P_i + c_N} - e_i g. \qquad (29)$$

Combing Eq. 28 and Eq. 29 recovers the well-known relationship between the ribosome mass-fraction ($f_R$) and growth rate ($g$)[26]:

$$g = \frac{p_R f_R c_N}{P_R + c_N}. \qquad (30)$$

The model also imposes additional consumption of the final metabolite product, $c_N$. Defining protein synthesis kinetics as $\rho_i = \frac{p_i(f_i r)c_N}{P_i + c_N}$ and using $\alpha_i = \frac{k_i E_i c_i}{K_i + c_i}$, we obtain

$$\frac{dc_N}{dt} = \alpha_{N-1} - \alpha_N - \rho_R - \sum_i^N \rho_i - c_N g.$$

Furthermore, we can derive the solution for steady-state growth rate following the same procedure as for Eq. 9, using $\sum_{i=1}^N \frac{dc_i}{dt} = 0$ and $\sum_{i=1}^N \frac{de_i}{dt} + \frac{dr}{dt} = 0$:

$$\sum_{i=1}^{N} \frac{dc_i}{dt} = \sum_{i=1}^{N-1} (\alpha_{i-1} - \alpha_i - gc_i) + \alpha_{N-1} - \alpha_N - \rho_R - \sum_{i=1}^{N} \rho_i - c_N g$$

$$= \alpha_0 - \alpha_N - g \sum_{i=1}^{N} c_i - \rho_R - \sum_{i=1}^{N} \rho_i = 0$$

and

$$\sum_{i=1}^N \frac{de_i}{dt} + \frac{dr}{dt} = \sum_{i=1}^N \rho_i - g \sum_{i=1}^N e_i + \rho_r - rg = 0.$$

Thus,

$$\sum_{i=1}^N \frac{dc_i}{dt} + \sum_i^N \frac{de_i}{dt} + \frac{dr}{dt} = \alpha_0 - g/\gamma_o - g \sum_{i=1}^N c_i - g \sum_{i=1}^N e_i - gr = 0,'$$

hence

$$g = \frac{\alpha_0}{\frac{1}{\gamma_o} + \sum_{i=1}^{N} c_i + \sum_{i=1}^{N} e_i + r}. \quad (31)$$

Eq. 31 describes a steady-state growth rate slightly modified from Eq. 9, in which the growth rate is further reduced by the concentration of enzymes (including ribosomes). Since the total protein concentration in an *E. coli* cell is ~7 mM while the total metabolite concentration is ~300 mM, the steady-state growth rate is impacted most strongly by metabolite concentrations and import ($\alpha_0$), thus making Eq. 31 approximately equivalent to Eq. 9.

In the model, we also assumed that the catalytic rate for protein synthesis (i.e., translational elongation rate) and the MM constant ($P_i$) are the same for each enzyme (including ribosomes). Using an average protein length in *E. coli* of ~330 amino acids (aa)[27] and an estimated *in vivo* translation elongation rate of 15–30 aa/s, we estimate that the catalytic rate for producing a single protein by a ribosome is 0.05–0.09 $s^{-1}$. The median enzyme catalytic rate in *E. coli* is ~20 $s^{-1}$ [9] (Fig. S15A), indicating that enzyme catalytic rates ($k_i$) are generally ~200-fold higher than the catalytic rate for protein synthesis. If we account for polysome behavior, in which a single transcript can be translated by ~10 ribosomes, the ratio of $k/\rho$ is ~20 (Fig. S15B). Furthermore, we assume that $P_i = K_i$ since the MM constant for translation is on the micromolar scale[24,25] similar to the majority of enzymes[21]. Finally, since our proteomics measurements indicate that $f_R = 0.35$ (Fig. 2) and $f_R + \sum_i^N f_i = 1$, we assume that $f_i = \frac{1-f_R}{N}$ for all *i*.

For a TSEN model with the same TSEN architecture as in Fig. 5C (5 reactions, 1 bottleneck), simulations including translation produced similar results to Fig. 5C upon a temperature shift from 27 °C to 37 °C, with a spike height of ~0.6 and an overall response time of ~2 doublings (Fig. S15C). Importantly, enzyme concentrations remained constant after the temperature upshift (Fig. S15D) and metabolite dynamics followed similar trajectories to the constant-enzyme model (Fig. 15E) and were ~10-fold larger than enzyme concentrations (Fig. S15D,E), as expected. Additionally, increasing the MM constant for translation decreased the growth rate and response time, while increasing the spike height (Fig. S15F). Interestingly, these predictions match the behavior observed experimentally under fusidic acid treatment or deletion of *tusA*/*tusB* (Fig. S11F,G), suggesting that the mechanism underlying these perturbations involves inhibition of binding to translational substrate targets (e.g., mRNA, tRNA).
